# Kinase–phosphatase cycles drive self-sustained oscillations of protein condensates

**DOI:** 10.64898/2026.10.27.762139

**Authors:** Emanuele Zippo, Le Qiao, Dorothee Dormann, Thomas Speck, Friederike Schmid, Lukas S. Stelzl

**Affiliations:** Institute of Physics, Johannes Gutenberg University Mainz, Mainz, Germany; Institute of Molecular Physiology, Johannes Gutenberg University Mainz, Mainz, Germany; Institute of Molecular Biology (IMB), Mainz, Germany; Institute for Theoretical Physics IV, University of Stuttgart, Stuttgart, Germany

## Abstract

Phase-separated protein condensates organize biochemical processes in cells. ATP-driven enzymatic reactions can change the physical properties and even trigger the dissolution of such condensates. Here we use molecular simulations to show how ATP-driven kinase and phosphatase reactions modulate the organization of TDP-43 low complexity domain (LCD) condensates. TDP-43 phosphorylation is enhanced by condensation, as CK1*δ* preferentially binds to and acts in TDP-43 LCD condensates, whereas dephosphorylation by the phosphatase calcineurin occurs predominantly in the dilute phase. This differential recruitment of kinases and phosphatases, together with the slow redistribution of kinases into newly formed condensates, triggers oscillatory appearance and disappearance of TDP-43 LCD condensates. The oscillations are self-sustained, persisting at constant chemical driving, showing that an antagonistic kinase–phosphatase pair can endow a condensate with autonomous dynamics rather than only setting its structure.

## Introduction

In cells, a myriad of biochemical reactions and molecular processes occur simultaneously and must be controlled in time and space^1^. Cells are therefore intricately and dynamically organized. Biochemical regulation and compartmentalization are provided by membrane-bound organelles, but can also arise from the interactions of proteins and nucleic acids through binding and self-assembly, or through their phase-separation into biomolecular condensates^2,3^. These membrane-less compartments provide distinct chemical environments in which metabolites^4^, ions^5^, nucleic acids and proteins are enriched or depleted^6^, and can thus facilitate assembly processes, speed up enzymatic reactions, but also sequester molecules. How biological sequences determine specificity, phase behavior and co-condensation is increasingly being understood through a combination of experiments and molecular dynamics (MD) simulations^7,8^. These phenomena have largely been rationalized using equilibrium concepts, such as interaction valence, solvent quality and surface tension.

Cells, however, fundamentally rely on chemical-fuel-driven non-equilibrium processes. Driven systems can sustain states and dynamics that are inaccessible in equilibrium^9,10^. For example, dissipation of a chemical fuel can drive the growth and maintenance of synthetic fibrils and the formation of vacuolar and vesicular structures in peptide-based coacervates^11–16^. Theory and continuum simulation models demonstrate that active processes can control the size of phase-separated droplets and suppress Ostwald ripening^17–20^. In addition to chemical reactions, active transport can also lead to microphase separation into many small droplets rather than macroscopic phase separation^21^. Chemical reactions and the steady supply of a chemical fuel can also create interfacial currents^22,23^, switch the shapes of droplets and induce chaotic flow in emulsions of droplets^24–26^.

Oscillatory dynamics are another hallmark of non-equilibrium systems, and theory has predicted that chemical reactions coupled to phase separation can produce self-sustained cycles of droplet growth and dissolution^27^. Experiments with DNA-based coacervates showed that two competing enzymatic activities, DNA ligase and DNA nuclease, can drive cyclic changes in the DNA components of the condensates^28^. However, sustained autonomous oscillations were not achieved in this system. Oscillations have also been observed in synthetic chemical systems through the coupling of droplet growth to sedimentation under gravity^29^. Being able to engineer oscillatory dynamics of condensates would also open directions in synthetic biology and systems chemistry, such as timekeeping devices for synthetic cells and the periodic sequestration of intermediates in a reaction network. The difficulty of achieving self-sustained condensate oscillations in vitro highlights the need to better understand how biological sequences, molecular structures and the specificity they impart influence each other in non-equilibrium steady states^28^.

ATP-driven enzymatic phosphorylation provides a paradigmatic example of a non-equilibrium process in cells. Kinase and phosphatase cycles enable the relay of messages in cell signaling cascades^30^. Phosphorylation alters protein-protein interactions and, when it occurs in disordered regions, it often alters the propensity of proteins to phase separate^31^. Phosphorylation can modify both conformational ensembles and intermolecular interactions and thereby reorganize condensate structure^32^. Phosphatases are also an essential component of the cellular regulation of protein phosphorylation. By removing phosphate groups, they can counteract kinase activity and revert phosphorylation, and thus play a similarly important role in regulating protein-protein interactions and phase separation. This opposing action of kinases and phosphatases establishes phosphorylation–dephosphorylation cycles that can continuously dissipate chemical fuel and drive biomolecular systems away from equilibrium. Depending on the protein sequence and phosphorylation pattern, phosphorylation can either promote or inhibit condensation^33,34^. For example, phosphorylation has been shown to suppress condensation of TAR DNA-binding protein 43 (TDP-43)^35^, whereas it can enhance condensation in other systems^36,37^. TDP-43 phosphorylation is of great interest, as it is strongly associated with neurodegenerative diseases, for example amyotrophic lateral sclerosis (ALS) and frontotemporal dementia (FTD), and regulates the protein’s disease-linked condensation and aggregation behavior^34,38^. How kinase-phosphatase cycles control TDP-43 phosphorylation and condensation is still unknown.

Recently, it was shown that a protein condensate is dissolved ahead of cell division by the action of a kinase. After cell division, a phosphatase removes phosphate groups from the protein chains and the condensate re-appears^39,40^. In this case, however, the cycle follows the cell-cycle clock rather than generating its own rhythm. Whether opposing kinase and phosphatase activities acting on a condensate can instead generate sustained oscillations autonomously remains an open question. Sastre et al. point out that stress granules, P-granules, and Cajal bodies are enriched in kinases while the surrounding cytoplasm would have opposing phosphatase activities, which raises the question whether enzymatic phosphorylation and dephosphorylation cycles could shape the organization of condensates in cells and also trigger oscillatory dynamics^29^.

In general, sustained oscillations require negative feedback, a delay in that feedback, and sufficient non-linearity^41^. Phosphorylation by a kinase and dephosphorylation by a phosphatase together form a so-called futile cycle: ATP is consumed, while the protein returns to its initial state. When the enzymes approach saturation, such a cycle can respond to changes in enzyme activity in a strongly non-linear, switch-like manner^42^. Continuous chemical cycling and a non-linear response, however, do not necessarily produce oscillations. For example, the Michaelis–Menten approximation of a sequential, distributive dual-site phosphorylation cycle does not produce sustained oscillations^43^. Oscillators based on post-translational modification (PTM) therefore rely on an additional, slower process. In the cyanobacterial KaiABC clock, this process is the assembly of the Kai proteins into complexes of defined stoichiometry^44^, and theory shows more generally that self-assembly can supply the delay^45^. In the Min system, the slow process is instead the ATP-driven binding of proteins to the membrane and their release from it^46^. In these examples, an additional structure, such as a membrane or a protein assembly, contributes to the feedback and the delay. Whether the phase separation of the modified protein can itself provide these ingredients in a kinase–phosphatase cycle remains largely unexplored.

Here, we use MD simulations to investigate how the chemical-fuel-driven interplay between kinase-mediated phosphorylation and phosphatase-mediated dephosphorylation regulates the organization of protein condensates. We show how to simulate two different types of enzymes simultaneously while maintaining thermodynamic consistency. We follow the chemical modification of individual sites on TDP-43 low complexity domain (LCD) chains, mediated by the kinase casein kinase 1*δ* (CK1*δ*) and the phosphatase calcineurin^47,48^, to see how this affects their intermolecular interactions, phase behavior and enzyme localization. This molecular-level description allows us to resolve the dynamics of complex molecular systems of multiple enzymes and hundreds of proteins in a non-equilibrium steady state (NESS). We find that competition between kinases and phosphatases can lead to self-sustained cycles of condensate dissolution and reformation within a specific range of enzyme concentrations. CK1*δ* preferentially associates with the condensates, enhancing phosphorylation, whereas calcineurin predominantly remains in the dilute phase. This asymmetry, together with the chemical driving supplied by ATP, gives rise to oscillations. Building on the molecular simulations, we construct a continuum model that shows how the differential partitioning of kinase and phosphatase, together with the delayed redistribution of the kinase into newly formed condensates, is sufficient to generate sustained oscillations. The model further reveals how the enzyme abundance controls the oscillation period. Together, our multiscale modeling establishes a design principle for enzymatically driven oscillatory dynamics of condensates.

## Results

### Thermodynamically consistent simulations of kinase-phosphatase cycles

We model a complete phosphorylation-dephosphorylation cycle with kinase CK1*δ* and phosphatase calcineurin on the substrate protein TDP-43 LCD using a combination of residue-level coarsegrained MD simulations^49,50^ and Monte Carlo moves^51–53^. While the phosphorylation reaction (catalyzed by the kinase) is fueled by ATP, the dephosphorylation reaction (catalyzed by the phosphatase) is usually not driven, but out-of-equilibrium concentrations of phosphate groups in the environment can still occur. Here, we allow only serine residues to get phosphorylated and dephosphorylated. The reaction is implemented as a bead exchange between serines (Ser) and phosphoserines (pSer), which occurs through a Monte Carlo step whenever the Ser/pSer residue is in contact with the active site of the enzyme. The acceptance probability for the Monte Carlo move is

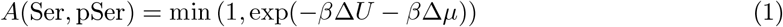

where *β* = 1/(*k*_*B*_*T* ), Δ*U* is the difference in potential energy between final and initial configurations and Δ*µ* is the chemical potential difference between thermostatted waste and fuel (respectively ADP + P_i_ and ATP, for phosphorylation reactions), fixed for the whole simulation. We will refer to the driving energy associated with phosphorylation reactions with Δ*µ*_k_ (related to kinase action) and to the one associated with dephosphorylation reactions with Δ*µ*_p_ (related to phosphatase action). Notice that these quantities are defined in such a way that the conversion of Ser into pSer is favored when Δ*µ*_*k,p*_ < 0 and, consequently, the dephosphorylation reaction is driven in forward direction when Δ*µ*_p_ > 0. Setting Δ*µ*_*k,p*_ to a value different from 0 breaks chemical equilibrium and detailed balance.

After checking through Markov state models (MSM) the thermodynamic consistency of both phosphorylation and dephosphorylation cycles with TDP-43 and either kinase or phosphatase (Supplementary Text, Supplementary Fig. 1, 2, 3, 4, 5, 6), we can build a complete phosphorylation-dephosphorylation cycle with kinase CK1*δ*, phosphatase calcineurin and substrate protein TDP-43 LCD (Methods, Supplementary Movie 1). We use as substrate a 5 times phosphorylated TDP-43 (5p TDP-43) to enhance the binding affinity and the frequency of reaction events.

We discretize the trajectory into 6 Markov states, as shown in Fig. 1a. The resulting MSM and the choice of the lag time for the estimate of the transition rates are validated by looking at implied timescales (Fig. 1b) and a Chapman-Kolmogorov test (Methods, Supplementary Fig. 7, 8). We repeated the simulations at drivings of Δ*µ*_k_ = 0, −5, −10 kJ mol^−1^, Δ*µ*_p_ = 0 kJ mol^−1^and also Δ*µ*_k_ = −5 kJ mol^−1^, Δ*µ*_p_ = +2.5 kJ mol^−1^. Fig. 1c shows the ratio of the forward and backward rates for each transition of 6-state MSM for each choice of the parameter pair (Δ*µ*_k_, Δ*µ*_p_). Again, the only transitions with broken detailed balance are the ones where the driving is applied (2 ↔ 3, 5↔ 6 for the case with Δ*µ*_k_ = − 5 kJ mol^−1^and Δ*µ*_p_ = +2.5 kJ mol^−1^). Also in this case, the heat dissipated in a complete phosphorylation-dephosphorylation cycle matches the sum of the driving of kinase and phosphatase (Fig. 1d).

**Figure 1:**
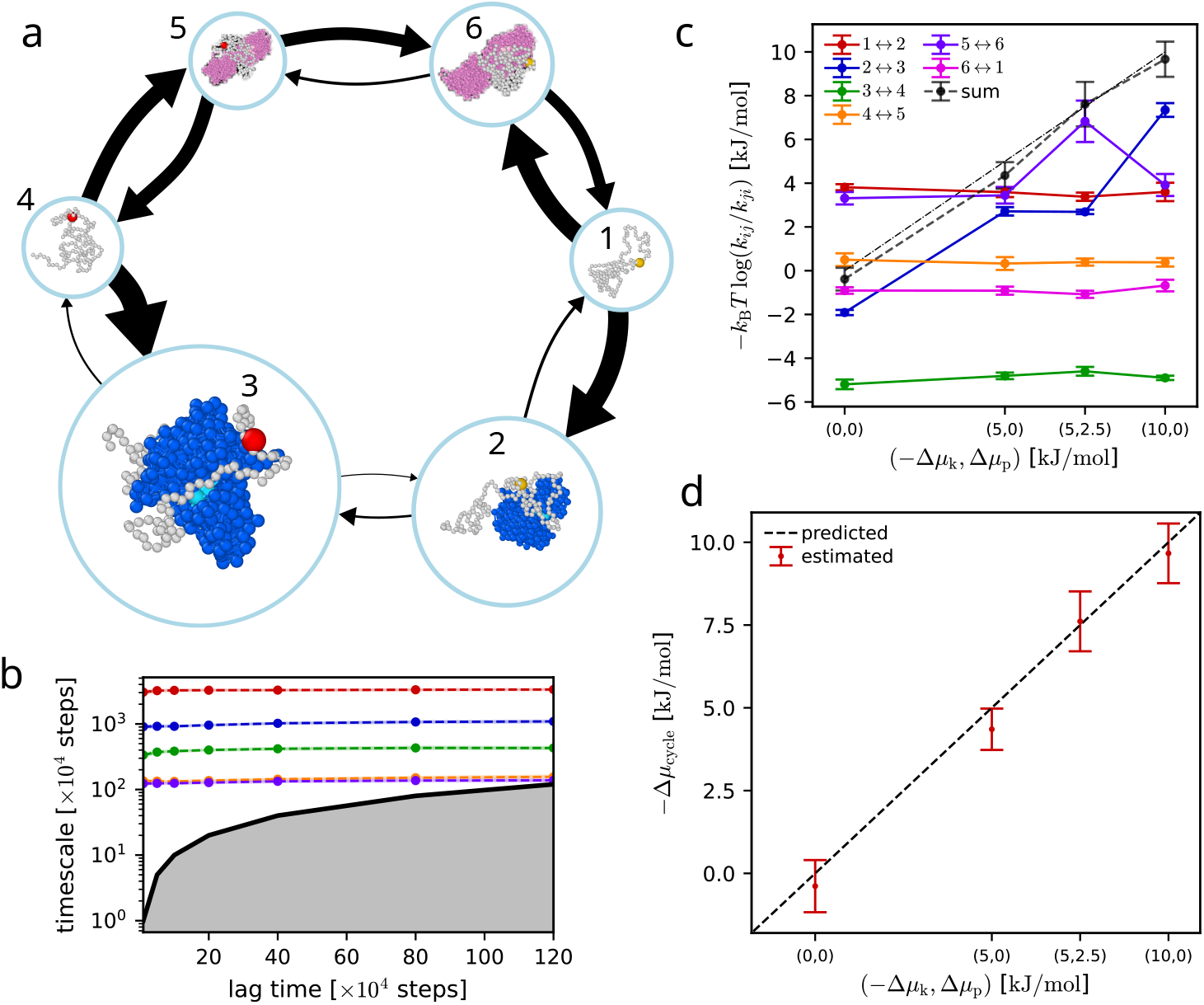
Thermodynamic consistency of complete phosphorylation-dephosphorylation cycles. (**a**) Sketch of the MSM for an example simulation with TDP-43 LCD (gray, pSer in red), CK1*δ* (blue) and calcineurin (magenta) at Δ*µ*_k_ = − 5 kJ mol^−1^ and Δ*µ*_p_ = 0 kJ mol^−1^. The size of the arrows is proportional to the absolute value of the corresponding transition rate and the size of the circular node is proportional to the stationary distribution of the corresponding state. (**b**) Implied timescales at different lag times from eigenvalues of the 6-state MSM transition matrix at Δ*µ*_k_ = − 5 kJ mol^−1^ and Δ*µ*_p_ = 0 kJ mol^−1^ (Methods). (**c**) Mean values +/-SEM of the single edge contributions to Δ*µ*_cycle_, i.e. *k*_*B*_*T* log(*k*_*ij*_/*k*_*ji*_). (**d**) Mean values +/-SEM of −Δ*µ*_cycle_ vs −Δ*µ*_k_ + Δ*µ*_p_.

Interestingly, state 3 in Fig. 1a is the most populated, followed by state 2, showing how TDP-43 binds preferentially to CK1*δ* than calcineurin, and that the binding affinity of CK1*δ* for more phosphorylated TDP-43 (6p TDP-43) is greater than for less phosphorylated TDP-43 (5p TDP-43). Calcineurin, instead, has a slight preference for the less phosphorylated TDP-43.

### Phosphatases counteract dissolution of hyperphosphorylated TDP-43 condensates

Given that hyperphosphorylation of TDP-43 LCD leads to condensate dissolution^35,52^, we asked whether adding phosphatases to hyperphosphorylated TDP-43 LCD in the homogeneous regime can trigger the reappearence of TDP-43 condensates. We tested this with simulations of 200 hyperphosphorylated TDP-43 LCD chains in the homogeneous phase and 5 calcineurin, with Δ*µ*_p_ = +8 kJ mol^−1^(Fig. 2a,b). To generate the starting hyperphosphorylated state, we performed simulations with TDP-43 and CK1*δ* at Δ*µ*_k_ = − 48 kJ mol^−1^(Methods). The kinases progressively hyperphosphorylate TDP-43 and eventually dissolve the condensate (Fig. 2c), in agreement with previous experiments^35^.

**Figure 2:**
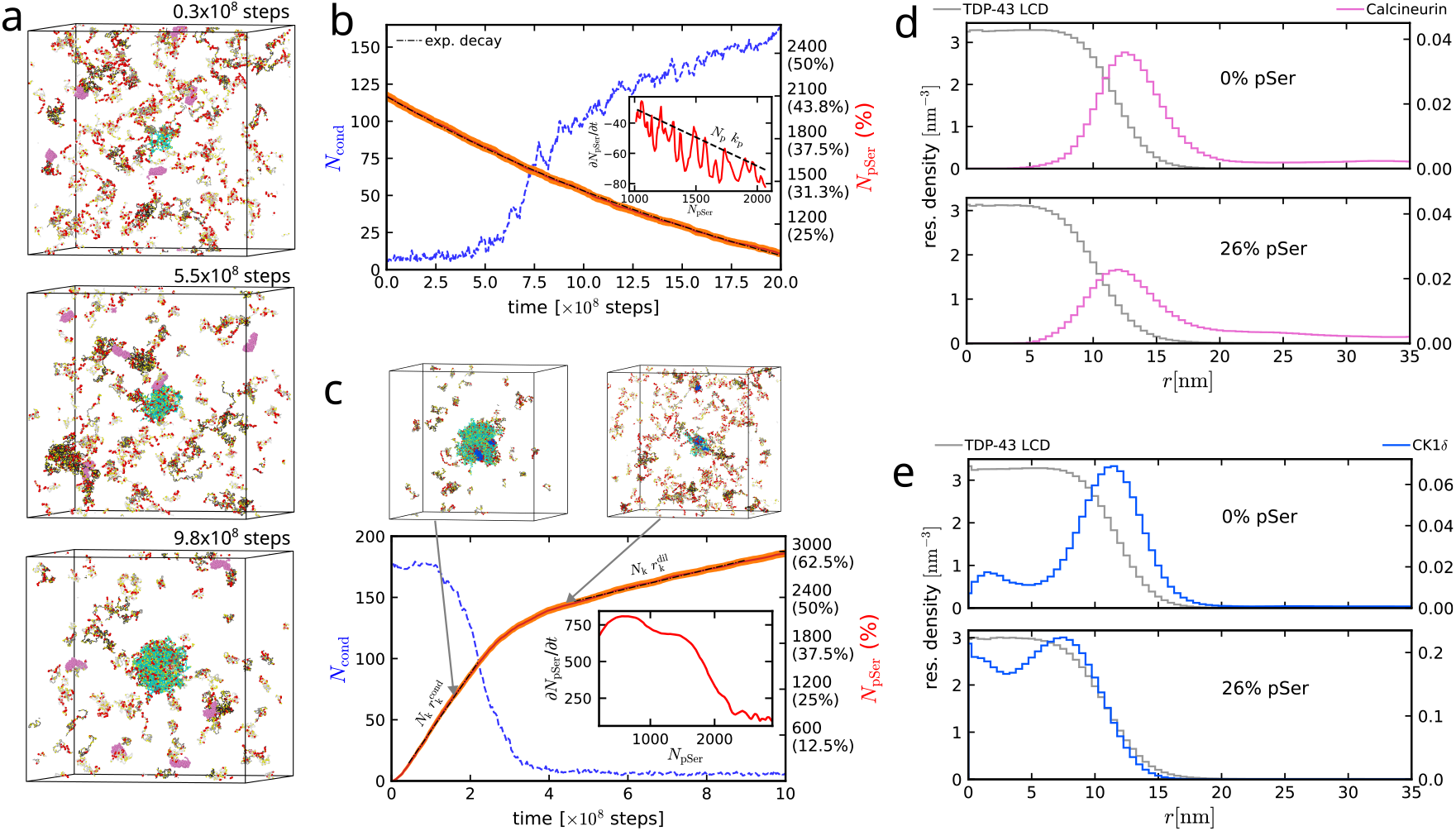
Enzymatic activity of calcineurin and CK1*δ* is determined by their partitioning in TDP-43 LCD condensate. (**a**) Snapshots from simulation with 5 calcineurin at times 0.3 *µ*s, 5.5 *µ*s and 9.8 *µ*s in simulation time, with biggest cluster (cyan) centered in the box. A TDP-43 condensate forms when the phosphorylation level drops. Calcineurin are in magenta and pSer in red. (**b-c**) Number of TDP-43 chains in condensate *N*_cond_ (blue, left y-axis) and total number of pSer *N*_pSer_ (+/-SEM, red, right y-axis, also in percentage over total number of Ser) as a function of time for simulations with 200 TDP-43 LCD and 5 calcineurin (b) or 5 CK1*δ* (c). The linear fits of log *N*_pSer_ (b) or *N*_pSer_ (c) in time, shown with dashed black lines, are used to extract decay rate *k*_p_ and phosphorylation rates in the phase-separated regime 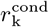 and in the homogeneous regime 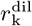. Insets show the numerical derivative of the red curves from linear fit with sliding interval of 21 points, plotted against *N*_pSer_. Boxes (c) show a snapshot in phase-separated regime and a snapshot in homogeneous regime. (**d-e**) Radial density profile of TDP-43 LCD (gray) and enzyme residues from equilibrium simulations with 5 calcineurin (d, magenta) or 5 CK1*δ* (e, blue). No TDP-43 Ser are phosphorylated in simulations for the upper panel, while 26% of them are phosphorylated for the lower panel. Counts are normalized by the volume of the spherical bin.

Calcineurin primarily interacts with hyperphosphorylated TDP-43 in the homogeneous aqueous regime. As more and more phosphate groups are removed from TDP-43 chains, a condensate starts to reappear (after about 6*µ*s in simulation time), when the number of chains in condensate *N*_cond_ rises above ∼50 and the number of phosphoserine *N*_pSer_ goes down to ∼1600, with a single chain having on average around 8 pSer (Fig. 2a,b, Supplementary Movie 2). The dephosphorylation dynamics is not strongly affected by the appearance of a condensate. We observe a first-order dynamics throughout the whole simulation time, with an exponential decay rate of the number of phosphates in time of *N*_p_*k*_p_ = −0.03802 ± 0.00004 *µ*s^−1^ (in simulation time), obtained from a linear fit of log *N*_pSer_ in time, where *N*_p_ is the amount of phosphatases. The inset in Fig. 2b shows the linear dependence of the time derivative of *N*_pSer_ on *N*_pSer_ itself. The dashed line shows the same line of best fit from the exponential decay. This behavior is explained by the fact that calcineurin is almost uniformly distributed in the box. Equilibrium simulations with unphosphorylated TDP-43 show that calcineurin mostly acts in the dilute phase and rarely binds to the condensate surface, while it does not enter the dense phase (Fig. 2d). At higher fraction of phosphorylated Ser residues, with condensate still mostly intact (e.g. at around 26% pSer), we observe a decrease in interaction between calcineurin and condensate, with an even lower percentage of phosphatase attached to the interface. As already observed from the MSM cycle (Fig. 1a, Supplementary Fig. 1), the interaction strength between calcineurin and TDP-43 is slightly higher for less phosphorylated chains compared to more phosphorylated ones. Indeed, calcineurin is overall negatively charged and the electrostatic repulsion lowers the total interaction strength with TDP-43 LCD chains (Supplementary Fig. 9). Interestingly, we also observe that the more phosphorylated C-termini of TDP-43 LCD chains (residue 361-414) preferentially locate at the interface of the condensate. By contrast, for the simulation with 0 pSer, C-termini and N-termini are equally distributed in the condensate (Supplementary Fig. 10).

In contrast to phosphatase, CK1*δ* activity is stronger in the phase-separated regime than in the homogeneous regime (Fig. 2c), confirming previous observations^52^. The average phosphorylation rate per kinase goes from 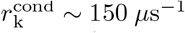 in condensate to 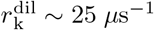 in the dilute phase (with time measured in simulation units). Since the kinase acts mostly on the condensate, where the density of substrate is high, its activity slows down when the condensate dissolves into a homogeneous solution, where the density of substrate is low. Moreover, we observe from equilibrium simulations at fixed amount of pSer that the kinase acts initially at the interface, but, as phosphorylation progresses, it can also enter the condensate (Fig. 2e). Indeed, CK1*δ* is positively charged, which results in a higher interaction strength between phosphorylated TDP-43 and kinases compared to unphosphorylated TDP-43 (Supplementary Fig. 9), as already shown by the MSM cycle in Fig. 1a. This behavior explains the initial acceleration of the phosphorylation rate between 0 and 0.2 × 10^8^ steps.

We verified the partitioning of the enzymes relative to the condensate through equilibrium simulations at fixed phosphorylation configurations using slab geometry and 2 different residuelevel force fields, namely CALVADOS3^49^ and HPS-Urry^50^ (Methods). To rule out finite size effects, we performed simulations with more TDP-43 LCD chains (400 chains). Slab simulations confirm that calcineurin locates in the dilute phase or at the interface of the condensate in the absence of phosphates, and show that an even smaller fraction of phosphatase interacts with the condensate in the case with 1715 pSer (18% pSer, Supplementary Fig. 11a,b). On the other hand, CK1*δ* has an opposite behavior: it mostly binds to the surface in the absence of phosphates and it also diffuses into the inside at 18% pSer (Supplementary Fig. 11c,d).

We also verified the partitioning of enzymes in condensates of full-length TDP-43. From slab simulations with 150 full length TDP-43 (both phosphorylated and not) and 5 enzymes (Methods), we see that calcineurin can move more easily inside the condensate and it is not repelled at the interface (Supplementary Fig. 12a,b). The condensate density is lower in this case, and the interaction strength with calcineurin is increased by the presence of the folded domains, particularly for the case with 922 pSer, allowing the phosphatase to move inside the dense phase (Supplementary Fig. 13). CK1*δ* is also more strongly sequestered in the full-length TDP-43 condensate compared to LCD (Supplementary Fig. 12c,d). The two force fields show overall a good agreement both for LCD and full-length TDP-43 condensates, with HPS-Urry hydropathy scale showing a somewhat stronger interaction between calcineurin and the condensate compared to CALVADOS3 and in general a stronger hydrophobicity of folded domains (Supplementary Text).

### The kinase-to-phosphatase ratio controls condensate phase behavior

At physiological ATP, ADP and P_i_ concentrations, phosphatase enzymes counteract the effect of adding kinases to TDP-43 condensates. We simulated 200 TDP-43 LCD in a cubic box with different numbers of CK1*δ* (*N*_k_) and calcineurin (*N*_p_) molecules and let the system reach a non-equilibrium steady state (NESS), where the activity of kinases and phosphatases is balanced and ATP is steadily consumed. Different concentration ratios between kinases and phosphatases give rise to different phosphorylation configurations in the steady state. We thus expect to be able to control the phase separation behavior by tuning enzyme concentration ratios while keeping the ATP, ADP and phosphate concentrations fixed.

As expected, increasing the number of phosphatases increases dephosphorylation activity and promotes condensation. For example, simulations with 2 kinases and 24 phosphatases form a stable condensate containing around 150 chains, whereas increasing the number of kinases to 4 and reducing the number of phosphatases to 8 leads to condensate dissolution (Fig. 3a,b). Small clusters can occasionally form due to thermal fluctuations, but rapidly disappear. The number of pSer fluctuates around 1000 residues for *N*_k_ = 2, *N*_p_ = 24, while it reaches approximately 2000 for *N*_k_ = 4, *N*_p_ = 8 (Fig. 3b, in red). Thus, the main determinant of condensate formation and size appears to be the number of phosphate groups in the system, which continuously changes due to enzymatic activity.

**Figure 3:**
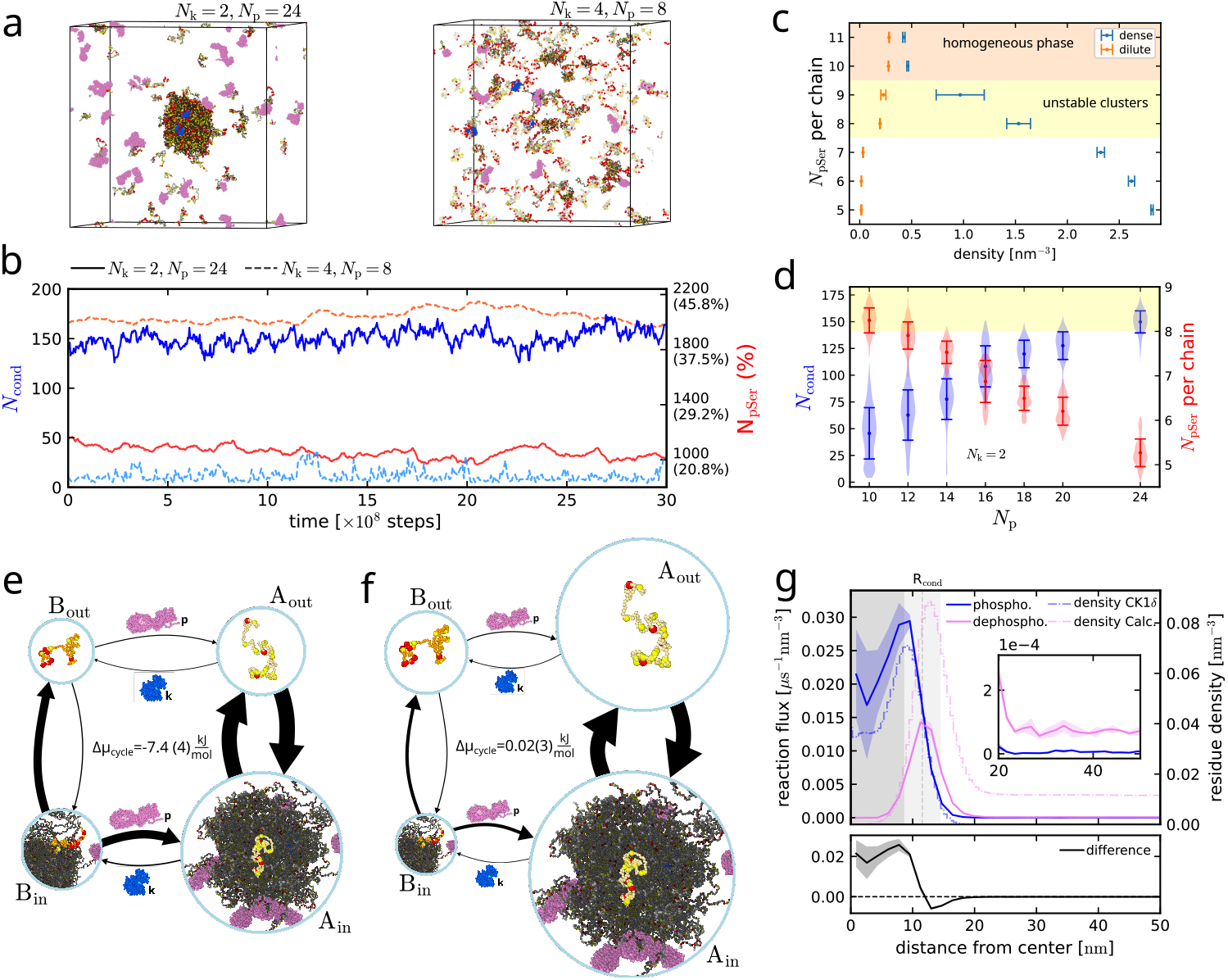
Phase separation propensity control through active tuning of phosphorylation state. (**a**) Snapshots from simulations at NESS with *N*_k_ = 2, *N*_p_ = 24 and *N*_k_ = 4, *N*_p_ = 8. Calcineurin is in magenta, CK1*δ* in blue and pSer in red. (**b**) Number of chains in condensate *N*_cond_ (blue) and total *N*_pSer_ (red, also in percentage) in time from simulations with 200 TDP-43 LCD and *N*_k_ = 2, *N*_p_ = 24 (solid lines) or *N*_k_ = 4, *N*_p_ = 8 (lighter dashed lines). (**c**) Phase diagram with *N*_pSer_ per chain vs TDP-43 bead density in dense (blue) and dilute phase (orange) from slab simulations without reactions. All TDP-43 have same number of pSer, but different phosphorylation patterns extracted at random from the simulations with 2 CK1*δ* and 24 calcineurin. Densities are extracted from fit of the density profile, errors are computed from the 3 different replicas. (**d**) Distributions with mean and standard deviation of *N*_cond_ (blue, left y-axis) and number of pSer per chain (red, right y-axis) for different amounts of calcineurin and 2 CK1*δ*, from 3 replicas. (**e,f** ) MSM cycle sketches showing transitions from chains of type A (low phosphorylation degree, yellow chain) to B (high phosphorylation degree, orange chain) and from dense to dilute phase, from simulations with 2 CK1*δ* and 24 calcineurin with (**e**, Δ*µ*_k_ = − 40 kJ mol^−1^, Δ*µ*_p_ = +8 kJ mol^−1^) and without driving (**f**, Δ*µ*_k_ = 0 kJ mol^−1^, Δ*µ*_p_ = 0 kJ mol^−1^). The size of the arrows is proportional to the absolute value of the corresponding transition rate and the size of the circular node is proportional to the stationary distribution of the corresponding state. (**g**) Phosphorylation (blue) and dephosphorylation (magenta) events per bin volume per *µ*s as a function of the distance from the center of the condensate (left y-axis) and residue density profile of CK1*δ* and calcineurin (respectively blue and magenta, dashed lines on right y-axis), from reactive simulation at *N*_k_ = 2, *N*_p_ = 24 with driving. Errors are computed from 3 replicas. The gray shaded area shows the dense phase, with vertical line at the condensate radius *R*_cond_ and light gray area of width 2*w*.

However, a universal phosphorylation threshold for phase separation is difficult to define because the phase behavior depends not only on the total number of phosphate groups, but also on their distribution among chains and their pattern along individual chains^32,54,55^. In order to characterize the phase-separation propensity of TDP-43 LCD as a function of phosphorylation degree, we simulate systems of 200 TDP-43 LCD in slab geometry, with a fixed number of pSer and without enzymes. The phosphorylation patterns are sampled from simulations with 2 CK1*δ* and 24 calcineurin, thereby averaging over the phosphorylation patterns observed in the reactive simulations while retaining their relative occurrence. By constructing a phase diagram of the number of pSer per chain versus the concentrations of dilute and dense phases (Fig. 3c, Supplementary Text), we find that chains with 7 pSer or less undergo phase separation, chains with 8 and 9 pSer form unstable clusters, and chains with 10 or more pSer remain in a homogeneous state.

Having established this equilibrium reference for the dependence of phase separation on phospho-rylation degree, we next fix the number of kinases to *N*_k_ = 2 and vary the number of phosphatases *N*_p_ from 10 to 24. Increasing *N*_p_ leads to a monotonic increase in the mean droplet size and decrease in the number of phosphate groups in the system at NESS (Fig. 3d). The distributions show distinct behavior at different enzyme concentrations. For *N*_p_ = 16, a bimodal distribution may result from insufficient time for the relaxation to a NESS or from differences in phosphorylation patterns and condensate composition between replicas. Comparison of the three replicas shows that the second peak at lower pSer is mainly caused by replica 2 (Supplementary Fig. 14a). Indeed, the phosphorylation pattern in replica 2 differs slightly from the others: Ser332 and Ser333 in the conserved region (residues 321–340) are more frequently phosphorylated, whereas other serines are less phosphorylated (Supplementary Fig. 14b). Instead, simulations at *N*_p_ = 10 and 12 show bimodal distributions of *N*_cond_ and *N*_pSer_ in all replicas (Supplementary Fig. 15a). Here, the smaller condensate-size peak corresponds to the dissolved state, which occurs when the average number of pSer per chain exceeds ∼ 8, consistent with the unstable clusters regime observed in the equilibrium simulations.

### Activity generates fluxes across the condensate interface

Although the propensity of each chain species to phase separate is an equilibrium property, enzymatic activity continuously shifts the system towards stronger or weaker phase separation. Kinases and phosphatases in competition generate a flux of chains between condensed and dilute phases: hyperphosphorylated chains leave the condensate and eventually encounter a phosphatase, which removes phosphate groups and restores their propensity to phase separate, as illustrated in Fig. 3e,f. To quantify the strength of the activity, we classify chains into two species according to their phos-phorylation state: species A comprises chains with 7 or fewer pSer and is prone to phase separation; species B comprises chains with 8 or more pSer and has a reduced propensity to phase separate. We then discretize the trajectories of individual TDP-43 chains from simulations with 2 CK1*δ* and 24 calcineurin into four states: A in the dilute phase, A in the dense phase, B in the dense phase, and B in the dilute phase. The A-to-B transition in the dense phase is predominantly driven by CK1*δ*, whereas the B-to-A transition in the dilute phase is predominantly driven by calcineurin.

From the ratio of clockwise to counter-clockwise transition probabilities, we estimate the average heat dissipated per cycle (Methods), analogous to our thermodynamic validation. The enzymatic activity generates a net clockwise flux corresponding to an average dissipation of Δ*µ*_cycle_ = 7.4± 0.4 kJ mol^−1^per chain cycle. In contrast, when the chemical driving is set to zero, the net cycle flux vanishes and the estimated dissipation per cycle is Δ*µ*_cycle_ = 0.02 ± 0.03 kJ mol^−1^.

Interestingly, the transition rate from B to A is also high in the dense phase. Indeed, many dephosphorylation reactions occur at the interface of the condensate, where calcineurin and pSer accumulate (Fig. 3g). By analyzing phosphorylation and dephosphorylation reaction fluxes as a function of distance from the condensate center, we find that phosphorylation dominates in the condensate core, whereas dephosphorylation dominates in the dilute phase. Both reactions contribute at the interface, with phosphorylation prevailing in its inner region and dephosphorylation in its outer region. This spatial organization generates a net chemical flux across the interface, consistent with previous theoretical studies^22,23^, and can be largely attributed to the spatial distribution of the enzymes relative to the condensate (Fig. 3g).

To resolve phosphorylation–dephosphorylation cycles at the interface in greater detail, we extended the MSM analysis to simulations saved at a higher temporal resolution, with a frame interval of 0.5 ns, and subdivided the dense-phase states into core and interface states (Methods). The interface width *w* = 2.93± 0.02 nm was determined from the density profile of TDP-43 LCD chains in driven simulations at NESS (Fig. 3g, shaded area). The resulting 6-state MSM confirms that most of the transitions between A and B occur at the interface (Supplementary Fig. 16) and reveals cycles between condensate core and interface, with an estimated dissipation Of 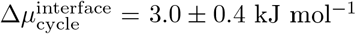, as well as cycles between the interface and dilute phase, with an estimated dissipation of 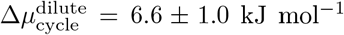 (Supplementary Fig. 17). Given the coarse-grained definition of species A and B, we cannot resolve the full dissipation, since most of the reactions happen within single states. Again, the analysis of the simulations with no driving results in no dissipation at the interface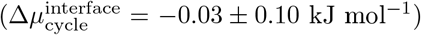. Indeed, reactions are in detailed balance and kinases and phosphatases can both phosphorylate and dephosphorylate chains, leading them to the equilibrium concentrations. Since calcineurin has access to phosphosites that are rarely phosphorylated by CK1*δ*, the resulting phosphorylation configurations are different from the active case (Supplementary Fig. 18a) and the system evolves towards a new configuration at around 5.5 pSer per chain and smaller condensate (around 110 chains in condensate) (Supplementary Fig. 18b).

### Predicting Ostwald ripening from molecular-scale kinetics

The discretization of the system into phase-separating and phase-separation deficient chains (A and B) and the computation of the MSM transition rates (Fig. 3e) allow us to predict from molecular scale whether droplets coarsen or can coexist at a selected size. Continuum theories of active emulsions of ternary fluids^9^ predict that Ostwald ripening is suppressed and multiple droplets can coexist if the radius of the stable condensates *R*_cond_ exceeds the threshold radius *R*_stab_. By relating the theoretical framework^56^ to our MD simulations, we can estimate *R*_stab_ as:

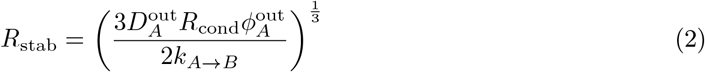

where 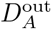 is the diffusion coefficient of the phase-separating species A in the dilute phase, 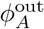 is its volume fraction in the dilute phase and *k*_*A→B*_ the rate of conversion of phase-separating chains into soluble chains.

For the system with *N*_k_ = 2, *N*_p_ = 24 at Δ*µ*_k_ = − 40 kJ mol^−1^, Δ*µ*_p_ = +8 kJ mol^−1^, we obtain a threshold radius of *R*_stab_ = 15.3 ± 0.3 nm (Methods), larger than the mean condensate radius of *R*_cond_ = 11.56 ± 0.05 nm. We thus expect to observe Ostwald ripening for this system, despite the enzymatic activity. Coexistence of multiple droplets could potentially be achieved by increasing the conversion rate from A to B in the dense phase by approximately threefold, for example, by increasing the enzyme concentrations while keeping the kinase-to-phosphatase ratio fixed. Despite the relatively fast phosphorylation rate, due to the kinase enrichment in the condensate, the net conversion of A chains into B is hindered by the activity of phosphatase at the interface, which recycles droplet material before the B chains can leave the condensate. On the other hand, simply increasing the droplet size by increasing the total amount of TDP-43 chains might not suppress Ostwald ripening. *R*_stab_ depends itself on *R*_cond_, and changing the size of the system might only impact *R*_stab_ indirectly through the other quantities. Moreover, this estimate assumes that the loss of droplet material scales with the droplet volume. Given the kinase activity at the interface and the slow redistribution of kinases from the dilute phase, this assumption may depend on the condensate composition and not hold for this system, potentially leading to even stronger Ostwald ripening.

To test this prediction, we doubled the simulation box in one direction and prepared a configuration with 2 droplets of different size, one in each half of the box, using configurations extracted from the previous simulation at NESS. We observe the larger droplet growing at the expense of the smaller one (Supplementary Movie 3). Simulations without reactions coarsen in the same way (Supplementary Fig. 19). The droplet size here is limited by the amount of phase-separation-competent material available, rather than selected by a balance of reaction and diffusion fluxes, despite the continuous influx and efflux of condensate material.

### ATP-driven kinase–phosphatase competition produces self-sustained oscillations

Oscillations are a hallmark of driven non-equilibrium system, but designing self-sustained oscillations in condensates has proven hard^28^, and we wondered whether our simulations could reveal design principles for self-sustained oscillations. At enzymes concentrations for which condensates become small and unstable (e.g. simulations with 2 CK1*δ* and 10 calcineurin, Fig. 3c,d), kinase activity can be switched off by the condensate dissolution. In contrast, phosphatase activity remains active, and can even increase slightly in the homogeneous phase, where a larger fraction of the hyperphosphorylated chains is accessible (see Fig. 2b,c). In this regime, dephosphorylation becomes dominant and dephosphorylated chains can nucleate a new condensate, where kinases will eventually dock. This non-linearity of the kinase activity, coupled with the time delay due to kinases diffusion into the new condensate, leads to alternating condensate dissolution and reformation (Fig. 4a,b, Supplementary Movie 4,5).

**Figure 4:**
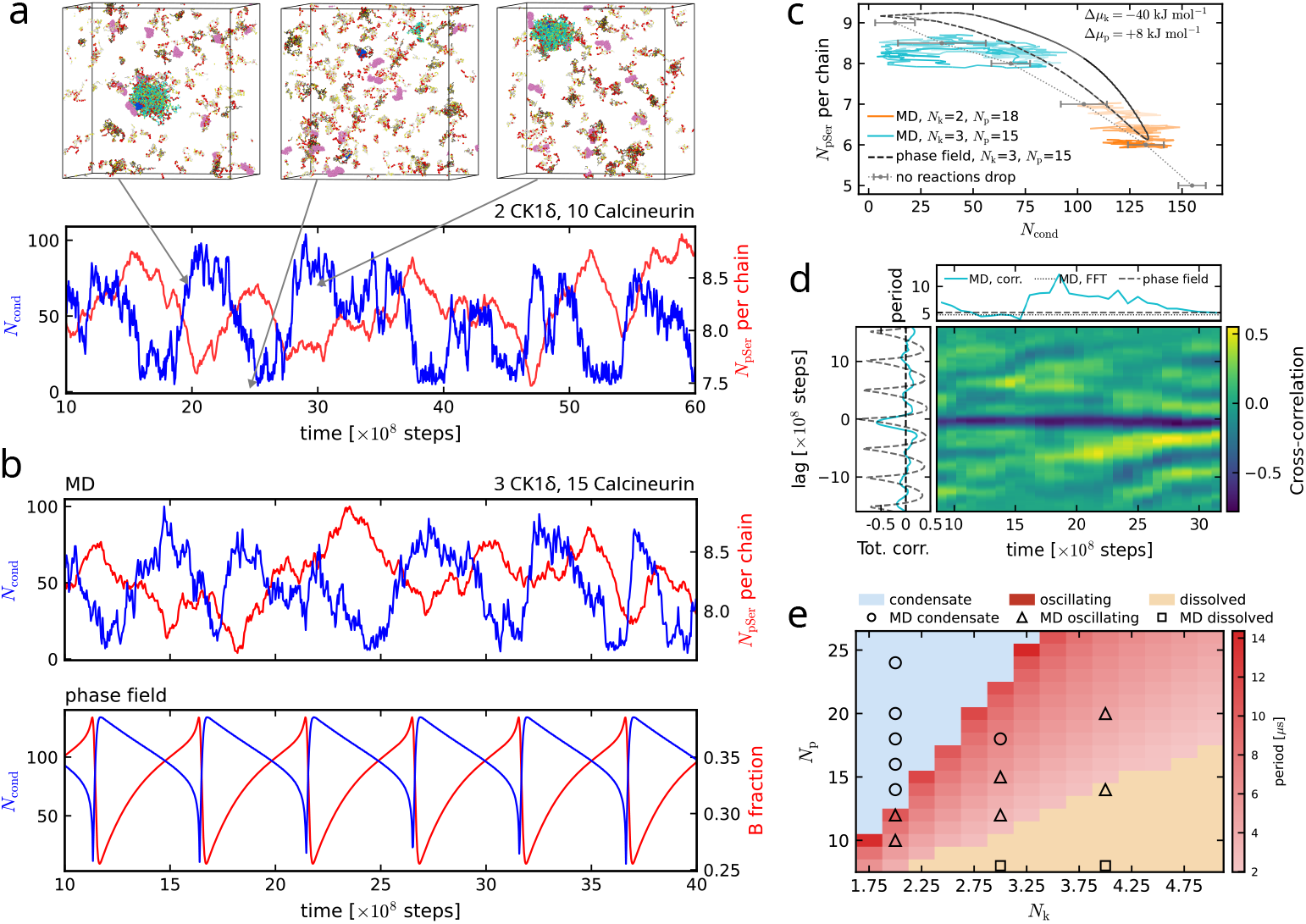
Kinase-phosphatase competition leads to oscillatory behavior in MD simulations and phase-field simulations. **(a)** Number of chains in condensate *N*_cond_ (blue, left y-axis) and *N*_pSer_ per chain (red, right y-axis) in MD trajectory with *N*_k_ = 2, *N*_p_ = 10. Insets show snapshots from MD simulation, with condensate disappearing and re-appearing. **(b)** *N*_cond_ (blue, left y-axis) for MD trajectory (up) and phase-field model (bottom) with *N*_k_ = 3, *N*_p_ = 15 (b). In red on the right y-axis *N*_pSer_ per chain for the MD trajectory and fraction *ϕ*_B_/(*ϕ*_A_ + *ϕ*_B_) for phase-field simulations. **(c)** Phase portrait with *N*_cond_ on x-axis and *N*_pSer_ per chain in y-axis with *N*_k_ = 2, *N*_p_ = 18 and *N*_k_ = 3, *N*_p_ = 15. The black line shows the phase portrait from phase-field simulations at *N*_k_ = 3, *N*_p_ = 15. Gray dashed line and points from equilibrium simulations with fixed amount of pSer, each point is the average of 3 replicas starting from different phosphates configuration with same total amount of pSer. **(d)** Cross-correlation between *N*_pSer_ and *N*_cond_ from MD simulation and phase-field simulations at *N*_k_ = 3, *N*_p_ = 15 (left). Pseudocolor plot of cross-correlation with sliding time window of 16 *µ*s from MD trajectory. On top oscillation period from cross-correlation at each window, computed as the interval between the 2 peaks closest to 0 lag (solid cyan line), from peak of the Fourier spectrum of the total cross-correlation (dotted black line) and from phasefield simulations (dashed gray line). **(e)** Phase diagram from phase-field simulations. Red: oscillating, colored by the period; pale blue: stable condensate; beige: dissolved. MD simulations: stable condensate (circles), oscillating (triangles), no condensate (squares).

For simulations at *N*_k_ = 2, *N*_p_ = 10 and *N*_k_ = 3, *N*_p_ = 15, we observe that the amount of phosphorylation oscillates around 8-9 pSer per chain, around the critical value for phase separation (Fig. 3c). When this value exceeds ∼ 8.25, the condensate dissolves and consequently the amount of phosphorylation starts to drop. After the condensate reforms, the kinases need time to diffuse into the newly formed condensate (Supplementary Fig. 20). We visualize this coupled dynamics through phase portraits of phosphorylation degree versus condensate size (Fig. 4c). For the oscillating simulation at *N*_k_ = 3, *N*_p_ = 15 (same as in Fig. 4b), the trajectory forms a closed loop characteristic of oscillating systems, whereas the stable-condensate simulation at *N*_k_ = 2, *N*_p_ = 18 relaxes towards a fixed point. We also report the phase portraits for all the other setups at *N*_k_ = 2 in Supplementary Fig. 21a. To provide an equilibrium reference, we compare these trajectories with simulations in equilibrium without enzymatic reactions at fixed phosphorylation degrees, initialized from configurations sampled from the reactive simulations. In the trajectory with oscillations, the system repeatedly overshoots the equilibrium phase boundary, driving the condensate towards smaller sizes and eventually dissolution, before dephosphorylation restores the conditions for condensation.

Starting from configurations at NESS, when we switch off the driving (Δ*µ*_k_ = 0 kJ mol^−1^and Δ*µ*_p_ = 0 kJ mol^−1^), oscillations stop. In particular, the system evolves towards a new configuration, as already observed for the case with stable condensate at *N*_k_ = 2 and *N*_p_ = 24 (Supplementary Fig. 18b). We repeated the procedure for *N*_k_ = 2 and *N*_p_ = 10 and 14. All simulations evolve towards the same fixed point in the phase diagram, at around 5.5 pSer per chain and 100-110 chains in condensate, assuming similar phosphorylation patterns (Supplementary Fig. 21b).

The oscillatory behavior of the system becomes even clearer when inspecting the cross-correlation between phosphorylation degree and cluster size. For the trajectories that present oscillations, we computed both the total cross-correlation and a time-dependent cross-correlation, with sliding window of 16 *µ*s in simulation time (Supplementary Fig. 22). As expected, we see strong anti-correlation near zero lag, confirming that phosphorylation follows condensate formation with little delay. We estimated the period of oscillation both directly from the time-dependent cross-correlation, as the time interval between the 2 peaks closest to 0 lag, and from the peak of Fourier power spectrum of the total detrended cross-correlation (Supplementary Fig. 23). For the example simulation with *N*_k_ = 3, *N*_p_ = 15, we observe sustained oscillations throughout the full trajectory, with oscillation period around ∼ 5*µ*s in simulation time (Fig. 4d). However, the power spectrum is, in general, broad and the total cross-correlations can be irregular and decay with time (Supplementary Fig. 24). This behavior may reflect both the limited simulation time and temporary losses of periodicity.

Such variations in periodicity may arise from stochastic fluctuations and rearrangements of phosphorylation patterns. For example, in the trajectory shown in Fig. 4a (*N*_k_ = 2, *N*_p_ = 10), the phosphorylation patterns differ between the first two oscillations and the third, where N-terminal serines are more phosphorylated and C-terminal serines are less phosphorylated, with the exception of Ser410 (Supplementary Fig. 15b). The time-dependent cross-correlations indicate that these interruptions in periodicity are temporary rather than evidence of globally damped oscillations in the trajectories shown in Supplementary Fig. 22. For example, at *N*_k_ = 2, *N*_p_ = 10, periodicity is temporarily lost around 32*µ*s, but recovers after approximately 5*µ*s. In this case, the condensate persists longer during one cycle due to the random dissociation of one kinase (Supplementary Fig. 20).

By contrast, simulations with higher phosphatase concentrations, in which the condensate does not dissolve, exhibit substantially more irregular cross-correlations (Supplementary Fig. 25). Interestingly, some replicas in this regime also show features suggestive of oscillatory behavior, including strong anti-correlation close to lag zero (e.g. *N*_k_ = 2, *N*_p_ = 14, or *N*_k_ = 3, *N*_p_ = 18, Supplementary Fig. 25). However, transient overshoots in the number of pSer or droplet size might randomly occur as the system relaxes towards a fixed point (Supplementary Fig. 26).

To identify the minimal mechanism responsible for the oscillations and to explore a broader range of enzyme concentrations, we developed a continuum phase-field model for the phase separation of phosphorylated TDP-43 LCD chains with enzymatic activity. We represent TDP-43 using two components, *ϕ*_A_ and *ϕ*_B_, corresponding to weakly and strongly phosphorylated, i.e., phase separating and non-phase separating chains, respectively, together with kinase and phosphatase fields, *ϕ*_K_ and *ϕ*_P_, in the solvent *ϕ*_s_ (Methods, Eqs. (12) and (13)). A reaction term proportional to *ϕ*_K_ in the transport equations can convert *ϕ*_A_ into *ϕ*_B_, while a term proportional to *ϕ*_P_ converts *ϕ*_B_ into *ϕ*_A_, with drivings Δ*µ*_k,p_ analogous to the Monte Carlo move in the MD simulations. Phosphatases are preferentially excluded from the dense phase, while kinases are attracted to it, with stronger attraction to *ϕ*_B_ (strongly phosphorylated TDP-43 is more attracted to CK1*δ*). We show in Supplementary Fig. 27 how *ϕ*_P_ diffuses out of the condensate after it forms, while *ϕ*_K_ moves towards it.

We mapped the kinase and phosphatase densities to the numbers of enzyme molecules, *N*_k_ and *N*_p_, and compared the results with the MD simulations. The phase-field model reproduces oscillations in five of the six MD simulations that display oscillatory behavior (Supplementary Fig. 22). A comparison for the case with *N*_k_ = 3, *N*_p_ = 15 is shown in Fig. 4b as an example. The model reveals the origin of the time delay: after the kinase-rich condensate dissolves, a new condensate forms away from the kinase, and the kinase must subsequently diffuse into the newly formed condensate before phosphorylation can resume (as also shown by the density profile in Supplementary Fig. 27). Without driving, the condensate formed by the same model is stable and does not oscillate (Supplementary Fig. 28). The phase portraits show stable limit cycles. For *N*_k,p_ at which we observe oscillations in MD simulations, we verified that the limit cycles do not depend on the initial conditions (Supplementary Fig. 29), including the case at *N*_k_ = 3, *N*_p_ = 15, shown in Fig. 4c. Here we plot on the y-axis the fraction of B times the total number of Ser in a TDP-43 LCD chain (24× *ϕ*_B_/(*ϕ*_A_ + *ϕ*_B_)) for comparison with *N*_pSer_ per chain. The mean phosphorylation (or B fraction) is mapped onto the MD simulations through the Flory–Huggins interaction parameter between A and solvent *χ*_As_, which places condensate dissolution at around 0.33–0.4 of B fraction, corresponding to 8–10 pSer per chain.

The cross-correlation between phosphorylation fraction and condensate size for *N*_k_ = 3, *N*_p_ = 15 (Fig. 4d) matches the oscillation period of the corresponding MD simulation, about 5 *µ*s. Cross-correlation comparisons for the other oscillating MD trajectories are shown in Supplementary Fig. 22.

Fig. 4e shows the state diagram of the phase-field model together with all available MD simulations. We identify three regimes: phase-separating with high amount of phosphatases, homogeneous with low amount of phosphatases, and an intermediate oscillating regime. The model reproduces 12 of the 14 simulated setups. Since the model does not include thermal noise, the oscillations from phase-field simulations are regular, in contrast to the MD simulations. Thus each setup has a well-defined period: 6.6, 10.4, 3.1, 5.0 and 4.6 *µ*s for (*N*_k_, *N*_p_) = (2, 10), (2, 12), (3, 12), (3, 15) and (4, 20), within the 3–10 *µ*s range of the MD simulations (Supplementary Fig. 22). At fixed kinase-to-phosphatase ratio, adding enzyme shortens the cycle (6.6, 5.0 and 4.6 *µ*s at (2,10), (3,15), (4,20)); at fixed kinase number, adding phosphatase lengthens it (2.6 to 11.0 *µ*s as *N*_p_ goes from 11 to 22 at *N*_k_ = 3).

In conclusion, our results show that the coupling between phosphorylation-dependent phase behavior, differential enzyme partitioning and delayed kinase recruitment can generate self-sustained condensate oscillations in a non-equilibrium steady state.

## Discussion

Previous theoretical work^9,19,20,24,27,56,57^ and experiments^14,16,18,29^ have established that chemical reactions can control both the size and dynamics of biomolecular condensates, but how this control emerges from molecular-scale interactions remains poorly resolved. Here, using thermodynamically consistent^51,52,58^, transferable^49,50,59,60^ reactive molecular dynamics simulations, we show how the interplay between enzymatic phosphorylation and dephosphorylation can regulate condensate organization and phase separation in a biologically relevant system. MD simulations allow us to explore dynamic phosphorylation patterns and reveal complex features that cannot be captured by representing the system as a binary or ternary mixture of molecular species. By feeding this molecular information into a phase-field model, we reproduce the oscillatory behavior observed in the molecular simulations and identify the physical mechanism that gives rise to condensate dissolution and reformation within a specific range of kinase and phosphatase concentrations.

Markov state modeling has emerged as a useful tool for studying enzyme kinetics and selfassembly^61–63^. Here, we employed MSM both to assess the thermodynamic consistency of the reactive simulations and to extract molecular information from condensate and single-chain simulations. The latter analysis revealed that highly phosphorylated TDP-43 has a stronger affinity for CK1*δ* and a weaker affinity for calcineurin than weakly phosphorylated TDP-43. These differences in binding affinity translate into distinct partitioning of the two enzymes between phosphorylated and unphosphorylated condensates. The resulting enzymatic activity therefore cannot be fully described by simple Michaelis–Menten kinetics: the effective interaction between enzyme and substrate evolves with the phosphorylation state of the substrate, its phosphorylation pattern and its propensity to phase separate. Thus, molecular recognition and phase separation are dynamically coupled. In particular, the difference in kinase activity between the dense and dilute phases is a key ingredient of the observed oscillatory behavior.

Oscillations of chemically driven droplets^27^ have been realized in synthetic systems, where a carbodiimide-fueled coacervate oscillates for hundreds of cycles^29^. There, however, the non-linearity and the delay are supplied by a nucleation threshold together with size-dependent sedimentation and fusion^29^. Enzymatic control of coacervates has been demonstrated with a ligase–nuclease pair acting on DNA, which produces cyclic but not self-sustained dynamics^28^, and in cells, where a kinase dissolves the pyrenoid condensate ahead of division, but the kinase activity follows the cellcycle clock rather than generating its own rhythm^40^. Known post-translational oscillators obtain their slow step from an ordered assembly or from cooperative binding to a membrane surface^44–46^. In the system described here, both the non-linearity and the delay are consequences of the phase separation itself, although through different mechanisms: the saturation concentration depends sharply on the phosphorylation state of the chains, and the kinase partitions into the phase it dissolves while the phosphatase acts in the dilute phase. This mechanism is consistent with the arrangement proposed on general grounds for biomolecular condensates^27,29^, which we realize here with a kinase–phosphatase pair and resolve at the level of individual modification events. Importantly, our molecular dynamics simulations suggest that the time required for kinases to partition into newly formed condensates sets the delay (Supplementary Fig. 20), a mechanism that is also captured by the continuum phase-field model. A similar spatial arrangement, with kinases enriched in the condensate and phosphatases acting predominantly in the surrounding solution^29^, has been reported for P granules^64^, stress granules^65,66^, and Cajal bodies^67^, which suggests that condensates in cells may possess the ingredients for autonomous cycling even where the observed dynamics follow an upstream clock. An intrinsic cycle of this kind need not compete with transcriptional or cell-cycle regulation; rather, it determines how a condensate responds to such regulation, setting the relaxation time, the sharpness of dissolution, and whether a slowly varying input is amplified or filtered. It is interesting to note that Hnisz et al. have also suggested that PTM-controlled clustering or phase separation mechanism could give rise to burst dynamics in the transcription of genes^68^.

Our results also suggest that it might be possible to rationally design protein condensate-based oscillators for synthetic biology and systems chemistry. By changing the number of enzymes, the phase of the oscillations can be influenced. For example, at fixed kinase concentration, by modifying the phosphatase concentration, we can tune the time fraction that the system spends in the dissolved state (e.g. with fewer phosphatases, the system spends more time in dissolved state and less in condensed state in one oscillation period). Within the oscillating region of parameter space, changing the enzyme concentrations also alters the period of the cycle. At fixed kinase-to-phosphatase ratio, increasing the total enzyme concentration increases the oscillation frequency. Phase-field simulations allow one to explore these dependencies over a broader parameter space than is accessible with molecular dynamics alone (Fig. 4e). This provides a framework for investigating how reaction kinetics and enzyme abundance can be used to tune condensate dynamics.

Phosphorylation fraction and condensate size provide useful coarse-grained descriptors of these dynamics, but the molecular simulations reveal a richer underlying state space. Chains with the same overall phosphorylation fraction can exhibit different phosphorylation patterns and conse-quently different phase-separation behavior (as shown in Supplementary Fig. 14). Chains can reorganize between successive oscillations, leading to different patterns and condensate compositions (Supplementary Fig. 15b). This heterogeneity is a plausible origin of the variability in oscillation period. It is invisible to a two-species description, and is one aspect in which particle-based simulations contribute beyond continuum models.

Beyond oscillatory dynamics, the molecular simulations suggest that enzymatic activity may also promote the formation of finite-sized structures. We observe that phosphorylated C-terminal regions of TDP-43 preferentially accumulate near the condensate interface, consistent with recent experimental and theoretical observations^8^ of micellar structures. Other kinds of microphases might also emerge from the enzymatic activity by suppression of Ostwald ripening^9,17,56^. We constructed an MSM, which shows how the activity of the enzymes creates a constant outward flux of hyper-phosphorylated material leaving the condensate, and an inward flux of dephosphorylated material re-entering the dense phase. In our system, phosphorylation controls the condensation propensity of the proteins and the droplet size is limited by the finite amount of material. However, suppression of Ostwald ripening through activity might be possible, for example, by increasing the conversion of condensate material into soluble chains inside the dense phase (Eq. (2), Methods). Such internally driven material exchange provides a possible molecular mechanism for maintaining non-equilibrium finite-sized structures.

## Methods

### Coarse-grained molecular dynamics simulations

Molecular dynamics simulations were run using the Python package HOOMD-blue v3.8.1^69^ and the residue-level CALVADOS3 simulation model^49,60,70^. HPS-Urry model^32,50,59,71^ was also employed in slab simulations for comparison. Following the prescription for CALVADOS3, simulations were run using the Langevin integrator with integration step of 0.01 ps and damping rate of 0.01 ps^−1^. The temperature for all CALVADOS3 simulations (except for the thermodynamic validation simulations) was set at 250 K to match the ratio with the critical temperature (T/T_c_) of the HPS-cationpi simulations from our previous work^52^. Instead, HPS-Urry slab simulations were performed at 300 K.

Protein residues are represented with single beads and solvent and ions are treated implicitly in the interaction potentials. Different amino acids are distinguished by Lennard-Jones diameter *σ*, charge *q*, mass *m* and hydropathy scale parameter *λ* (parameters for CALVADOS3 simulations from Cao et al.^49^ and for HPS-Urry simulations from Regy et al.^50^). Consecutive beads are connected through harmonic springs with elastic constant of 8033 kJ mol^−1^nm^−2^(and 8360 kJ mol^−1^nm^−2^for HPS-Urry simulations). For residues in disordered regions, beads are placed at the C_*α*_ and the equilibrium distance is fixed at *r*_0_ = 0.381 nm. Instead, for residues in folded regions, beads are placed at their center-of-mass, with equilibrium distance which depends on the underlying structure. In order to keep the structure stable, elastic network was used between beads within each folded region at distances below 0.9 nm, with elastic constant 700 kJ mol^−1^nm^−2^. The folded structure of CK1*δ* was provided by PDB 6RU7 (X-ray crystallography)^72^. We used residues from 3 to 294 for our simulations. Instead, calcineurin is a complex of two proteins, which we connected through the elastic network as a single body. We used the structure provided by PDB 6NUF (X-ray crystallography)^73^.

Salt-screened electrostatic interactions between charged residues are modeled through the Debye-Hückel potential:

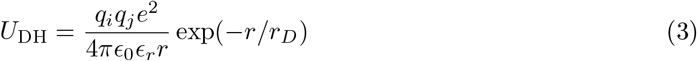

with *r*_*D*_ the Debye length, *e* the electron charge, *ϵ*_0_ the vacuum permittivity and *ϵ*_*r*_(*T* ) the temperature-dependent dielectric constant. The Debye length is computed as *r*_*D*_ = (8*πl*_*B*_*I*)^−1/2^, with ionic strength *I* = 150 mM and *l*_*B*_(*ϵ*_*r*_) the Bjerrum length. The Debye-Hückel interaction is cut off at distances above 4 nm (and 3.5 nm for HPS-Urry simulations).

Van der Waals interactions are modeled using the Ashbaugh-Hatch potential (a modified Lennard-Jones where attraction is scaled down based on the hydropathy of each specific amino acid pair):

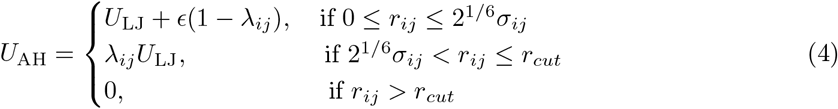

where *σ*_*ij*_ = (*σ*_*i*_ + *σ*_*j*_)/2, *λ*_*ij*_ = (*λ*_*i*_ + *λ*_*j*_)/2 and *U*_LJ_ = 4*ϵ* (*σ*_*ij*_/*r*_*ij*_)^12^ − (*σ*_*ij*_/*r*_*ij*_)^6^ is the standard Lennard-Jones potential, with *ϵ* = 0.8368 kJ mol^−1^and *r*_*cut*_ = 2 nm. Phosphorylated Ser residues are modeled as described by Rauh et al.^32,70^.

The box for simulations with single TDP-43 chain in the presence of enzymes is cubic with side length of 30 nm, while for condensate simulations the side length is 100 nm. We also employed slab geometry to check the position of the enzymes relative to the condensate. The box size used is 25 ×25 ×150nm^3^. All simulations employ periodic boundary conditions.

The Ashbaugh-Hatch pair potential for the non-bonded interactions is available at https://github.com/ezippo/ashbaugh_plugin as a HOOMD-blue plugin. The code used for the simulations is available at https://github.com/ezippo/hoomd3_phosphorylation.

### Phosphorylation and dephosphorylation reactions through Monte Carlo moves

We alternate MD steps with attempted Monte Carlo moves in order to mimic phosphorylation and dephosphorylation reactions. Phosphorylation reactions are implemented as:

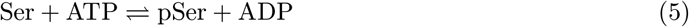

activated by the kinase CK1*δ* and with fixed implicit ATP/ADP concentration ratio parametrized by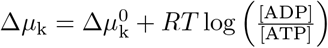. Instead, the dephosphorylation reaction reads:

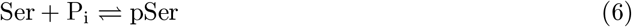

activated by the phosphatase calcineurin. This reaction is not fueled by ATP, but a possible imbalance of P_i_ can be modeled implicitly through a parameter 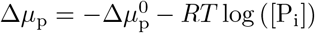. The sign convention is chosen such that the forward reaction is always in direction Ser →pSer. In the phosphorylation reaction, the presence of ATP will make Δ*µ*_k_ < 0 and push the reaction forward, while the dephosphorylation reaction will have Δ*µ*_p_ > 0 in order to push the reaction backward (pSer → Ser). Notice that 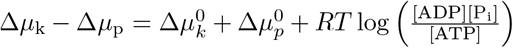is the energy released in an ATP hydrolysis reaction.

During the simulation, every 200 MD step, we check if the enzymes active site form a contact with a Ser or pSer bead. A contact occurs when the distances between all the residues belonging to the predefined active site and the Ser/pSer bead are below 1.05 nm. We chose as CK1*δ* active site the area close to Asp149, Phe150, Gly151 and for calcineurin the area close to Asp92, His94, Asp120, Asn152, His283. If a contact is detected, Ser and pSer beads are swapped with probability defined in Eq. (1) (Metropolis move^74^). The probability is biased by the chemical potential parameter Δ*µ*, which represents Δ*µ*_k_ in the case of a phosphorylation reaction (catalyzed by CK1*δ*) and Δ*µ*_p_ in the case of a dephosphorylation reaction (catalyzed by calcineurin). After accepting a Monte Carlo attempt to change particle identity from Ser to pSer or vice versa, new velocities for this particle are drawn from a Maxwell-Boltzmann distribution.

### Thermodynamic consistency through Markov-state modeling

The thermodynamic consistency procedure was applied to simulations with single TDP-43 with one reactive Ser, kinase CK1*δ* and phosphatase calcineurin. Following the procedure depicted in a previous publication^52^, we discretize the simulation trajectories into Markov states, we compute the dissipated heat per cycle from the transition rates, and we compare the result with the driving energy Δ*µ*. In order to optimize the estimation of the MSM, we lowered the temperature to 200 K and used as substrate TDP-43 LCD with 5 fixed phosphorylated serines among the most reactive (Ser369, Ser387, Ser393, Ser409, Ser410) to enhance the binding with CK1*δ*. We chose Ser403 as reactive phosphosite, as one of the most reactive Ser residues. We ran 5 simulation replicas for 20 × 10^8^ MD steps (20 *µ*s in simulation time) and dumped snapshots every 5000 MD steps, which were used to determine the states of the discretized trajectory (Supplementary Movie 1).

The states are determined by combination of phosphorylation state of the reactive site (Ser or pSer) and binding to the enzymes (bound or unbound), resulting in 6 possible states: state 1 is unbound 5p TDP-43, state 2 is 5p TDP-43 bound to CK1*δ*, state 3 is 6p TDP-43 bound to CK1*δ*, state 4 is unbound 6p TDP-43, state 5 is 6p TDP-43 bound to calcineurin, state 6 is 5p TDP-43 bound to calcineurin (Fig. 1a). As a simplification, we introduce a WCA-like repulsion between the residues of kinase and phosphatase to avoid spurious states where TDP-43 is bound to both enzymes. We used a Lennard-Jones energy *ϵ* = *k*_*B*_*T*/100 and repulsion distance *σ* = 12 nm.

Enzyme and substrate are considered bound when the distance to the enzyme active site of the closest residue of the substrate is less than 3 nm (roughly the size of the enzyme), and unbound when it is greater than 8 nm. For distances between 3 and 8 nm we apply a transition-based state assignment filtering^75^. Transition rates *k*_*ij*_(*τ* ) are estimated through the Maximum Likelihood estimator and uncertainties through Bayesian sampling using the Python package Deeptime^76,77^. We chose a lag time *τ* = 70 MSM steps (3.5 ns in simulation time). At greater or equal lag times, the implied timescales, computed as − *τ/* log |*l*_*i*_|, with *l*_*i*_ the eigenvalues of the transition matrix, remain stable (Supplementary Fig. 8). We also checked the goodness of the estimated MSM through a Chapman-Kolmogorov test (CK test)^78,79^ by comparing *k*_*ij*_(*nτ* ) (the transition matrix estimated at lag *nτ* ) and *k*_*ij*_(*τ* ) ^*n*^ (the transition matrix predicted from the one at lag *τ* ) for every possible transition *i* ⇌ *j* and different *n* (Supplementary Fig. 7).

We computed the ratio of the transition rates for each branch of the MSM network from simulations at different driving Δ*µ*_k_, Δ*µ*_p_. We expect the transitions between bound and unbound states to be constant and determined by the difference in free energy between the final state *j* and initial state *i*:

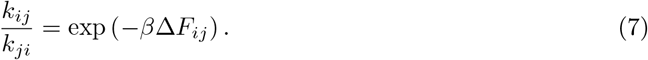

Instead, the transitions between phosphorylated and unphosphorylated states must depend on the driving:

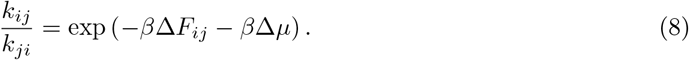

The sum of the contributions from all the branches of the MSM network has to match the total driving. Following this logic, we can directly compute the dissipated heat per cycle from the ratio between rates in clockwise direction and anti-clockwise direction, assuming that non-adjacent transitions are not allowed by construction (or very rare when the lag time is larger than 1 MD step):

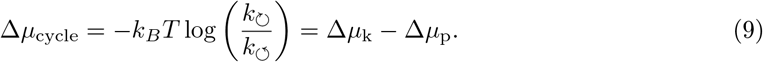

### Preparation of condensate simulations with reactive enzymes

We equilibrated a cubic box of 100 nm side length with 200 unphosphorylated TDP-43 LCD and inserted 5 CK1*δ* in the dilute phase at Δ*µ*_k_ = − 48 kJ mol^−1^, to mimic the driving from ATP hydrolysis. We let 4 simulation replicas evolve for 10 × 10^8^ MD steps (10 *µ*s in simulation time), till complete dissolution of the condensate (Fig. 2c).

We used these simulations as a starting point for the other setups. To simulate the re-condensation through phosphatases, we extracted snapshots at around 2100 pSer (when hyperphosphorylated TDP-43 LCD chains are in the homogeneous phase), removed CK1*δ* and added 5 calcineurin in the dilute phase at Δ*µ*_p_ = +8 kJ mol^−1^. Then we let the system evolve for 20 × 10^8^ MD steps (20 *µ*s in simulation time), till a condensate reappears (Fig. 2a,b, Supplementary Movie 2).

In order to find a NESS for simulations with different concentration ratios of kinases and phosphatases, we started again by taking TDP-43 configurations from snapshots of the dissolution simulations with 5 CK1*δ*. We run 3 simulation replicas with *N*_k_ = 2, *N*_p_ = 10 from snapshot at 2000 pSer for 25 × 10^8^ MD steps, with *N*_k_ = 2, *N*_p_ = 12 from snapshot at 1818 pSer for 20 × 10^8^ MD steps, with *N*_k_ = 2, *N*_p_ = 20, 24 from snapshot at ∼1670 pSer for 12.5 × 10^8^ and 11 × 10^8^ MD steps, with *N*_k_ = 4, *N*_p_ = 8 and 20 from snapshot at ∼1750 pSer for 12.5 ×10^8^ and 4.5 ×10^8^ MD steps respectively (Supplementary Fig. 30).

We then used equilibrated snapshots at NESS from these simulations to run all the simulations with competing enzymes shown in the main text, in order to save simulation time for equilibration. Setups with *N*_k_ = 2, *N*_p_ = 10, 12, 20, 24 and with *N*_k_ = 4, *N*_p_ = 8, 20 were continued and considered already equilibrated (unless explicitly stated in Supplementary Table 1). Setups with *N*_k_ = 2, *N*_p_ = 14, 16, 18 were started from snapshots extracted from *N*_k_ = 2, *N*_p_ = 20, setups with *N*_k_ = 3, *N*_p_ = 12, 15, 18 were started from snapshots extracted from *N*_k_ = 2, *N*_p_ = 10, while setups with *N*_k_ = 4, *N*_p_ = 14 were started from snapshots extracted from *N*_k_ = 4, *N*_p_ = 20 and subsequently *N*_k_ = 3, *N*_p_ = 8 were started from *N*_k_ = 4, *N*_p_ = 14.

**Table 1:**
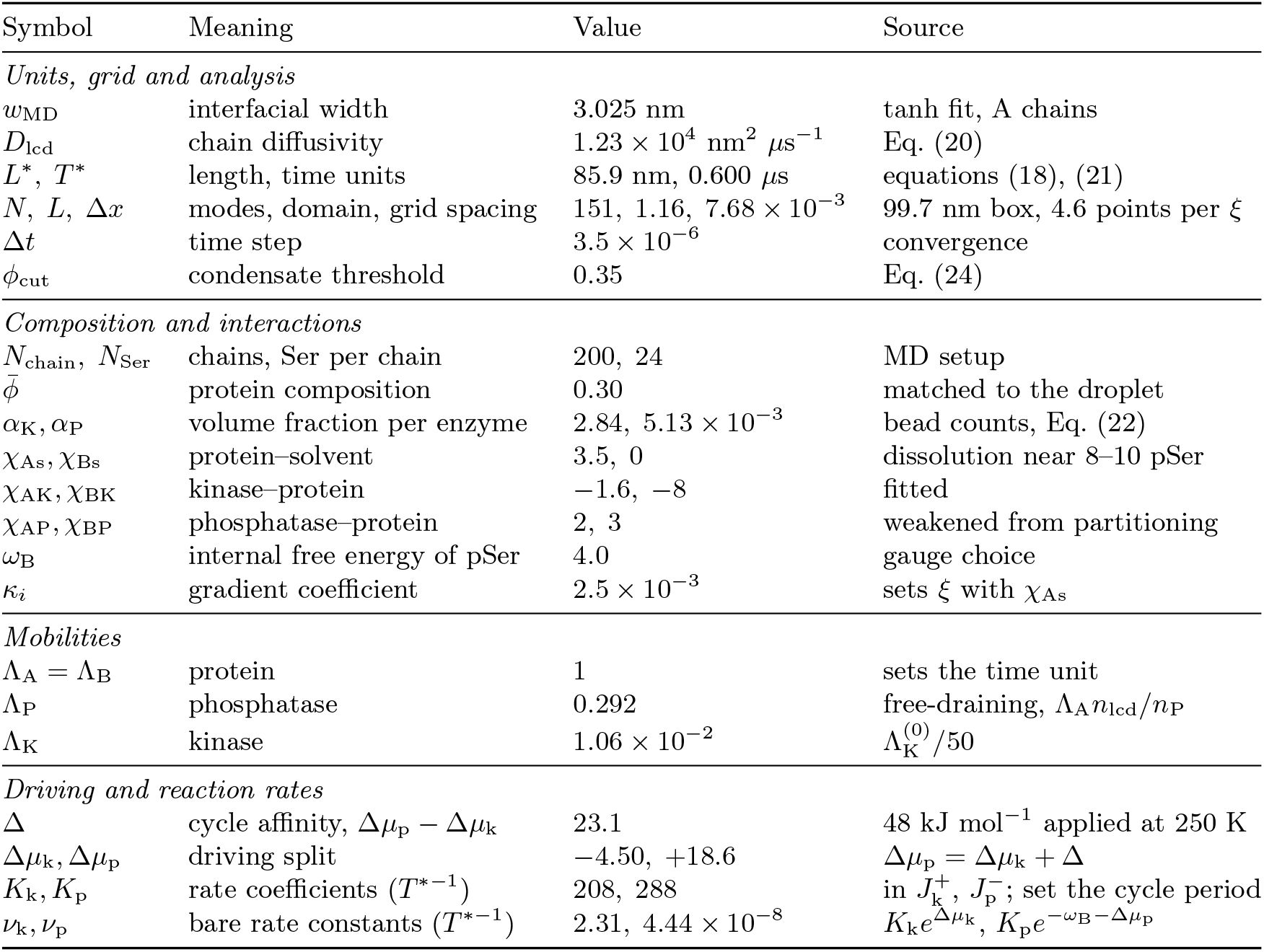
Continuum model parameters. Energies in *k*_B_*T* ; derivations are given in Methods.

| Symbol | Meaning | Value | Source |
|---|---|---|---|
| *Units, grid and analysis* |  |  |  |
| w<sub>MD</sub> | interfacial width | 3.025 nm | tanh fit, A chains |
| D<sub>lcd</sub> | chain diffusivity | 1.23 × 10<sup>4</sup> nm<sup>2</sup> µs<sup>−1</sup> | Eq. (20) |
| L<sup>*</sup>, T<sup>*</sup> | length, time units | 85.9 nm, 0.600 µs | equations (18), (21) |
| N, L, Δx | modes, domain, grid spacing | 151, 1.16, 7.68 × 10<sup>−3</sup> | 99.7 nm box, 4.6 points per ξ |
| Δt | time step | 3.5 × 10<sup>−6</sup> | convergence |
| φ<sub>cut</sub> | condensate threshold | 0.35 | Eq. (24) |
| *Composition and interactions* |  |  |  |
| N<sub>chain</sub>, N<sub>Ser</sub> | chains, Ser per chain | 200, 24 | MD setup |
| φ̄ | protein composition | 0.30 | matched to the droplet |
| α<sub>K</sub>, α<sub>P</sub> | volume fraction per enzyme | 2.84, 5.13 × 10<sup>−3</sup> | bead counts, Eq. (22) |
| χ<sub>As</sub>, χ<sub>Bs</sub> | protein–solvent | 3.5, 0 | dissolution near 8–10 pSer |
| χ<sub>AK</sub>, χ<sub>BK</sub> | kinase–protein | −1.6, −8 | fitted |
| χ<sub>AP</sub>, χ<sub>BP</sub> | phosphatase–protein | 2, 3 | weakened from partitioning |
| ω<sub>B</sub> | internal free energy of pSer | 4.0 | gauge choice |
| κ<sub>i</sub> | gradient coefficient | 2.5 × 10<sup>−3</sup> | sets ξ with χ<sub>As</sub> |
| *Mobilities* |  |  |  |
| Λ<sub>A</sub> = Λ<sub>B</sub> | protein | 1 | sets the time unit |
| Λ<sub>P</sub> | phosphatase | 0.292 | free-draining, Λ<sub>A</sub>n<sub>lcd</sub>/n<sub>P</sub> |
| Λ<sub>K</sub> | kinase | 1.06 × 10<sup>−2</sup> | Λ<sub>K</sub><sup>(0)</sup>/50 |
| *Driving and reaction rates* |  |  |  |
| Δ | cycle affinity, Δμ<sub>p</sub> − Δμ<sub>k</sub> | 23.1 | 48 kJ mol<sup>−1</sup> applied at 250 K |
| Δμ<sub>k</sub>, Δμ<sub>p</sub> | driving split | −4.50, +18.6 | Δμ<sub>p</sub> = Δμ<sub>k</sub> + Δ |
| K<sub>k</sub>, K<sub>p</sub> | rate coefficients (T<sup>*−1</sup>) | 208, 288 | in J<sub>k</sub><sup>+</sup>, J<sub>p</sub><sup>−</sup>; set the cycle period |
| ν<sub>k</sub>, ν<sub>p</sub> | bare rate constants (T<sup>*−1</sup>) | 2.31, 4.44 × 10<sup>−8</sup> | K<sub>k</sub>e<sup>Δμ<sub>k</sub></sup>, K<sub>p</sub>e<sup>−ω<sub>B</sub>−Δμ<sub>p</sub></sup> |

All information about the setups are listed in Supplementary Table 1.

### Preparation of slab simulations

In order to understand how enzymes interact with TDP-43 condensates, we simulated both TDP-43 LCD (400 chains) and full-length (150 chains) condensates, with either 5 CK1*δ* or 5 calcineurin, in slab configuration (Supplementary Fig. 11,12). For both cases we distributed the chains randomly in a large cubic box and shrank it to a side length of 25 nm to let the chains form a cluster. Then, we slowly extended the z axis to 150 nm and let the system thermalize.

Similarly, for the slab simulations used for the phase diagram in Fig. 3c, we shrank the large box to 22 nm and then elongated to 200 nm in z direction. In this case, we ran 3 different replicas with all chains having same number of pSer. We repeated for 5,6,7,8,9,10 and 11 pSer per chain. Phosphorylation patterns were extracted randomly from the 3 trajectories at *N*_k_ = 2,*N*_p_ = 24.

### Markov state model of condensate at NESS

We showed through MSM how the enzymatic activity creates a flux of material in and out of the condensate for simulations at *N*_k_ = 2, *N*_p_ = 24, for which we had a stable droplet. At each snapshot, we assigned each of the 200 TDP-43 LCD chains either to condensate or to dilute phase, using the connectivity-based clustering method from the Python package freud^80^ with 1 nm connectivity cutoff distance. We assumed that chains do not enter and exit the condensate with a frequency which is higher than our snapshot dumping frequency. By means of the record of reactions generated from our simulations code, we assigned each chain either to type A (with 7 or less pSer, phase separation prone) or B (with 8 or more pSer, phase separation deficient) and built an exact chemical timeline for each chain. Merging phase timelines and chemical timelines for each chain, we built a discretized trajectory of 4 states: 1 type A in dilute regime, 2 type A in dense regime, 3 type B in dense regime, 4 type B in dilute regime.

Given the chain-level discretized trajectories, we then computed the transition rates between the 4 states as the counts of transitions from *i* to *j*, divided by the total residence time in state *i*^81^. Errors are computed as square root of the counts from *i* to *j*, divided by the total residence time in state *i*. The dissipated heat per cycle is computed from the local detailed balance condition, using again Eq. (9).

Analogously, we built a 6 state MSM where we divided the states inside the condensate between core and interface regions. For this purpose, we used 3 new replicas of 10^9^ steps (10 *µ*s in simulation time) with finer dumping rate of 50 000 steps (0.5 ns). At each snapshot, we determined the condensate radius *R*_cond_ from the 90th percentile of distances from the cluster center of the residues belonging to the cluster. The interface region was defined as the spherical shell with radius between *R*_cond_ − *w* and *R*_cond_ + *w*, with *w* = 2.93 ±0.02 nm the interface width, while the core region is the sphere of radius *R*_cond_ − *w*. The width *w* was determined by the hyperbolic tangent fit of the TDP-43 residue density profile (Fig. 3g, Supplementary Text). Chains of the cluster with at least 25% of residues in the interface region were assigned to the interface states, while the rest to the core states.

### Ostwald ripening prediction

The theory for an active emulsion of ternary fluids^9,56^, predicts the possibility of suppression of Ostwald ripening through activity in externally maintained droplets. The stability analysis of a system of multiple droplets at radius *R*_cond_, the stable radius for a single droplet system, predicts that coexistence and suppression of Ostwald ripening is possible only if the slowest rate of relaxation to steady state is negative, i.e. if *R*_cond_ >*R*_stab_, with *R*_stab_ computed as in Eq. (2).

We computed *R*_stab_ for the system with *N*_k_ = 2, *N*_p_ = 24 with driving. We considered species A chains (with fewer than 8 pSer per chain) as phase-separating and species B chains (with at least 8 pSer per chain) as soluble. The stable radius of the droplet *R*_cond_ was computed from the density profile of the active droplet simulations (Fig. 3g). 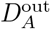 was extracted from the mean squared displacement (MSD) of the center of mass of A chains in dilute regime from the active droplet simulations. We computed the volume fraction of A in dilute regime as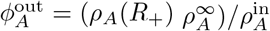, where 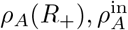 are the residue concentrations from the active droplet simulations computed just outside the interface (at the lowest density, before the supersaturation due to the phosphatases action far from the droplet kicks in) and in the dense phase, 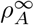 is the dilute phase residue concentrations from slab equilibrium simulations without curvature (see Supplementary Fig. 11). The rate of conversion of A into B was computed from the MSM in Fig. 3e as

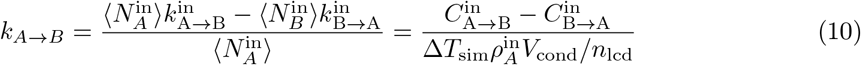

where *C*_*i→j*_ are transition counts from *i* to *j*, Δ*T*_sim_ is the total simulation time, *n*_lcd_ = 154 is the number of beads in one TDP-43 LCD chain and *V*_cond_ = ^4^ *πR*^3^ . In fact, 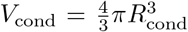 is equal to the concentration of A chains in condensate 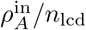 times the condensate volume *V*_cond_. In this way, we exclude the conversions from A to B in the dilute phase, not considered in the theory, and take into account possible recycling of droplet material A inside the dense phase by subtracting the counts of transition from B to A inside. Finally we compute *R*_stab_ as

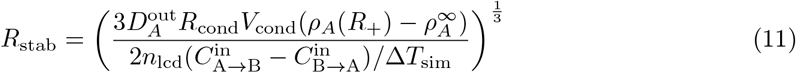

From the 3 replicas with *N*_k_ = 2, *N*_p_ = 24 at Δ*µ*_k_ = − 40 kJ mol^−1^, Δ*µ*_p_ = +8 kJ mol^−1^, we obtained *R*_stab_ = 15.3 ±0.3 nm, while *R*_cond_ = 11.56 ±0.05 nm<*R*_stab_. Thus we do not expect any suppression of Ostwald ripening in this case.

Consistently, passive simulations with *N*_k_ = 2, *N*_p_ = 24 at Δ*µ*_k_ = 0 kJ mol^−1^, Δ*µ*_p_ = 0 kJ mol^−1^lead to a vanishing net flux from A to B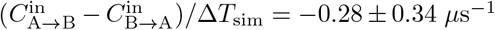, resulting in *R*_stab_ → ∞.

We tested the prediction of Ostwald ripening in our active emulsion system with simulations and compared it with passive simulations without reactions. We extracted 2 snapshots from the active simulations with *N*_k_ = 2, *N*_p_ = 24 with droplets of different size centered in the box, and placed them together in an elongated box with doubled side length in z direction, taking care of possible clashes at the boundaries (Supplementary Fig. 19a, Supplementary Movie 3). For 2 out of 4 replicas, we observed the bigger droplet growing at the expense of the smaller one within 5 *µ*s in simulation time. We also extended one of the replicas with 2 persisting droplets for 5 *µ*s for 3 independent runs. They all resulted in a single final droplet (Supplementary Fig. 19b,c). We repeated the same procedure, but switching off the reactions. We observed a similar behavior, as expected for passive systems (Supplementary Fig. 19d,e).

### Continuum phase-field model

To identify the physical ingredients underlying the oscillatory condensate dynamics observed in MD, we developed a coarse-grained phase-field model of TDP-43 in the presence of CK1*δ* and calcineurin. We describe the condensate by four volume-fraction fields on a one-dimensional periodic domain of length *L*: weakly phosphorylated protein *ϕ*_A_, heavily phosphorylated protein *ϕ*_B_, kinases *ϕ*_K_ and phosphatases *ϕ*_P_, with solvent *ϕ*_s_ = 1 −*ϕ*_A_ −*ϕ*_B_ −*ϕ*_K_−*ϕ*_P_. Transport conserves *ϕ*_A_, *ϕ*_B_, *ϕ*_K_, and *ϕ*_P_ separately, and the reactions interconvert *ϕ*_A_ and *ϕ*_B_. The local fraction of heavily phosphorylated protein is *p* = *ϕ*_B_/(*ϕ*_A_ + *ϕ*_B_). For comparison with MD, we use the effective mapping *N*_pSer_ = 24*p*, where 24 is the number of phosphorylation sites considered in the TDP-43 LCD.

The free energy is a Flory–Huggins mixture with square-gradient terms,

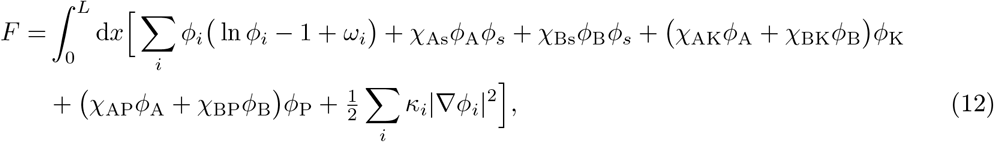

with *i* ∈{A, B, K, P, s} . Here *ω*_*i*_ is the internal free energy of species *i*, in units of *k*_B_*T* as are all energies of the model, with *ω*_A_ = *ω*_K_ = *ω*_P_ = *ω*_*s*_ = 0 fixing the zero baseline and *κ*_*s*_ = 0; *χ*_*ab*_ is the Flory–Huggins interaction parameter between species *a* and *b*, positive for repulsion; and *κ*_*i*_ is a square-gradient coefficient, with 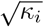 roughly of the order of the molecular size. The enzyme couplings determine the preferred spatial localization of the enzymes: *χ*_AK_, *χ*_BK_ < 0 attract the kinase to the protein, more strongly to phosphorylated protein, while *χ*_AP_, *χ*_BP_ > 0 exclude the phosphatase from the dense phase.

Writing the total protein as *ϕ* = *ϕ*_A_ + *ϕ*_B_, the two protein–solvent terms combine to *ϕ ϕ_s_χ*_eff_ (*p*) with *χ*_eff_ (*p*) = *χ*_As_ + (*χ*_Bs_ − *χ*_As_)*p*. Since *χ*_Bs_ <*χ*_As_, phosphorylation improves solvent quality and promotes dissolution. The difference *χ*_Bs_ − *χ*_As_ controls this contribution to the coupling between phosphorylation and phase behavior. Each species *i*∈{ A, B, K, P} moves down the gradient of its exchange chemical potential 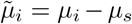, where *µ*_*i*_ = *δF/δϕ*_*i*_ is taken with all five volume fractions as independent variables. With mobility coefficient Λ_*i*_, the evolution equations are

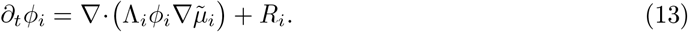

The first term describes diffusive transport, and *R*_*i*_ accounts for phosphorylation and dephosphorylation. Since the reactions interconvert A and B without consuming either enzyme, *R*_A_ = *R*_B_ and *R*_K_ = − *R*_P_ = 0.

The reactions are local. To better illustrate their dependence on composition, we split each chemical potential into ideal and excess parts, 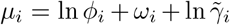, where 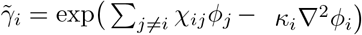 contains the interactions and the gradient contributions. The conversion A ⇌ B proceeds through two enzyme-catalyzed reactions, one catalyzed by the kinase and one by the phosphatase, labeled *c* ∈ {k, p}. The net rate of conversion of A into B is the sum over both reactions of a forward rate (A→B, phosphorylation) minus a backward rate (B→A)^10^,

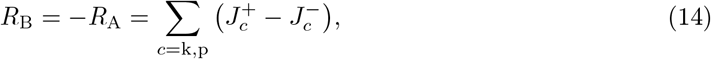

with the four rates

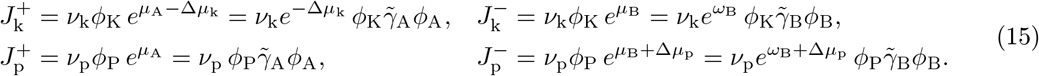

Here *ν*_k_ and *ν*_p_ are rate constants, and Δ*µ*_k_ and Δ*µ*_p_ create the chemical driving forces for the two reactions, with the same sign convention as in the MD simulations: ATP hydrolysis gives Δ*µ*_k_ < 0, which favors phosphorylation, and Δ*µ*_p_ > 0 favors dephosphorylation. Each rate is proportional to the local fraction of the enzyme and of the protein species it converts. In the limit of ideal gas chemical potentials, 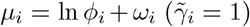, Eq. (15) therefore reduces to second-order mass-action kinetics, for instance 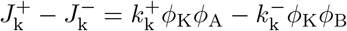 with constant coefficients 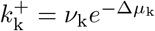 and 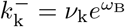. Interactions make the rate coefficients depend on the local composition through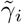.

The kinetics obey local detailed balance: for each reaction, the ratio of the forward to the backward rate is fixed by the free-energy change of converting one A into B in that reaction. From Eq. (15) and the definition of *µ*_*i*_,

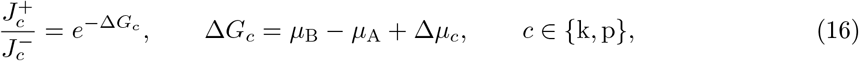

where *µ*_B_ − *µ*_A_ is the change in protein free energy and Δ*µ*_*c*_ the chemical free energy associated with reaction *c*. The enzyme fractions cancel from these ratios: the enzymes set how fast a reaction proceeds, but not in which direction. Each reaction therefore proceeds in the direction that lowers the free energy,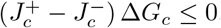.

This has two consequences. Without driving, i.e., at Δ*µ*_k_ = Δ*µ*_p_ = 0, both reactions push the protein towards chemical equilibrium, *µ*_A_ = *µ*_B_. If the system evolves according to Eq.(13), the free energy *F* of Eq. (12) then decreases monotonically, and the system relaxes to thermodynamic equilibrium; no sustained cycle is possible. In the presence of driving, phosphorylation of one A unit by the kinase followed by its dephosphorylation by the phosphatase forms a closed cycle, in which the kinase reaction runs forward and the phosphatase reaction backward. The free-energy change of this cycle is Δ*G*_k_− Δ*G*_p_ = Δ*µ*_k_ − Δ*µ*_p_ = − Δ. The protein chemical potentials cancel, so every cycle releases the same free energy Δ = Δ*µ*_p_− Δ*µ*_k_, corresponding to one hydrolyzed ATP, regardless of the local composition; this energy input sustains the cycle. We verified both properties numerically: without driving, *F* decreases at every time step, and with driving the heat dissipated per ATP consumed equals Δ to within 10^−4^. The driving applied in the MD simulations, Δ*µ*_k_ = − 40 kJ mol^−1^and Δ*µ*_p_ = +8 kJ mol^−1^, gives Δ = 48 kJ mol^−1^, or 23.1 *k*_B_*T* at 250 K.

In the general formulation above, some of the parameters are redundant: the transformation *ω*_B_ →*ω*_B_ + *δ*, Δ*µ*_k_ →Δ*µ*_k_ −*δ*, Δ*µ*_p_ →Δ*µ*_p_ −*δ, ν*_k_ →*ν*_k_*e*^−δ^ leaves every trajectory unchanged, because *ω*_B_ enters the dynamics only through the reaction rates; diffusion depends only on gradients of *µ*_B_. The kinetics thus depend on *ω*_B_, Δ*µ*_k_ and Δ*µ*_p_ only through Δ, the combination *ω*_B_ + Δ*µ*_k_, and the prefactors of the phosphorylation rate of the kinase and of the dephosphorylation rate of the phosphatase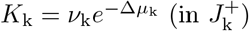 and 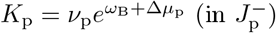.

The parameter *ω*_*B*_ +Δ*µ*_k_ quantifies the driving force and the reversibility of the kinase reaction: the backward rate coefficient of the kinase is 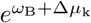 times its forward coefficient *K*_k_. *K*_k_ and *K*_p_ are second-order coefficients, setting the rate per unit enzyme and substrate volume fraction, and carry units of *T*^*−1^ because volume fractions are dimensionless. Because *ω*_A_ = 0 places the zero baseline of the internal free energy on the unphosphorylated species, *ω*_B_ enters *K*_p_, but not *K*_k_; the larger value of *K*_p_ therefore does not necessarily imply that the phosphatase is the more efficient enzyme.

The MD simulations contain *N*_chain_ = 200 TDP-43 LCD chains in a cubic box of side 100 nm. A chain is a string of *n*_lcd_ = 154 beads. The simulations further contain *N*_k_ CK1*δ* molecules (*n*_K_ = 292 beads) and *N*_p_ molecules of calcineurin (*n*_P_ = 527 beads). These two numbers of enzyme molecules are what the simulations varied, and they are the axes of Fig. 4e.

We determine the length scale from the interface of the unphosphorylated, enzyme-free binary reference system. Its bulk free-energy density, up to terms that do not affect coexistence, is

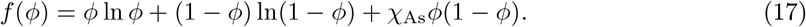

Let Δ*f* (*ϕ*) = *f* (*ϕ*) − *f*_tan_(*ϕ*) denote the excess above the common tangent connecting the coexistence compositions *ϕ*_dil_ and *ϕ*_dense_. The equilibrium planar interface obeys *κ*_A_(*∂*_*x*_*ϕ*)^2^/2 = Δ*f* (*ϕ*). Matching its midpoint slope to the profile *ϕ*(*x*) = *ϕ*_m_ + *A* tanh[(*x* − *x*_0_)*/ξ*], with *ϕ*_m_ = (*ϕ*_dense_ + *ϕ*_dil_)/2 and *A* = (*ϕ*_dense_ − *ϕ*_dil_)/2, gives

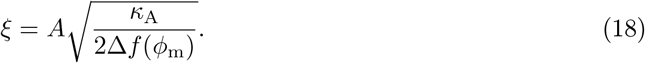

For *χ*_As_ = 3.5 and *κ*_A_ = 2.5 × 10^−3^, the coexistence fractions are *ϕ*_dil_ = 0.0379 and *ϕ*_dense_ = 0.962, giving *ξ* = 0.0352. Matching this width to the MD value *w*_MD_ = 3.025 nm yields

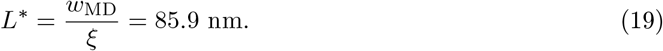

The time scale is calibrated using the dilute, free-draining diffusivity of an LCD chain. For independent Langevin friction on each bead,

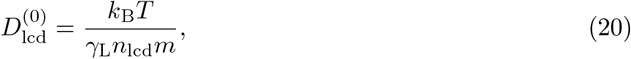

where *γ*_L_ = 0.01 ps^−1^is the damping rate and *m* = 110 Da is the mean bead mass. At *T* = 250 K, this estimate gives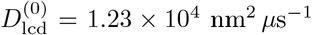. It is a free-draining reference value, rather than a measured diffusivity inside the condensate. In the dilute, long-wavelength limit, gradients of the exchange potential reduce to those of ln *ϕ*_*i*_, and Eq. (13) reduces to ordinary diffusion with physical diffusivity *D*_*i*_ = Λ_*i*_*L*^*2^/T^*^. Setting Λ_A_ = 1 and matching to 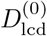 gives

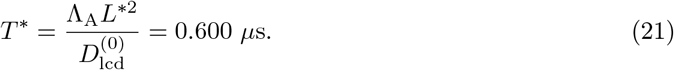

The continuum domain length *L* = 1.16 therefore corresponds to *LL*^*^ = 99.7 nm, approximately the side length of the MD box.

We set the mean total protein fraction to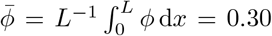. Assuming equal effective volume per MD bead, enzyme numbers are mapped to the spatially averaged enzyme fractions 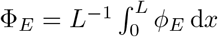 through

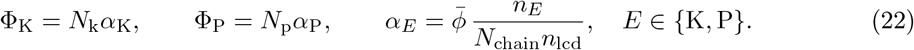

This matches enzyme-to-protein volume ratios using the chosen 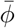. For freely diffusing enzymes, the same free-draining approximation gives

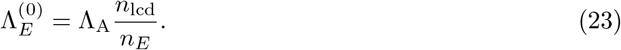

We use 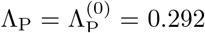 for calcineurin and a reduced effective kinase mobility 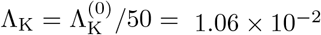, representing slow redistribution through substrate binding.

To quantify condensation, we define the region Ω_cond_(*t*) = {*x* : *ϕ*(*x, t*) >*ϕ*_cut_ }with *ϕ*_cut_ = 0.35. The integrated condensed protein fraction is

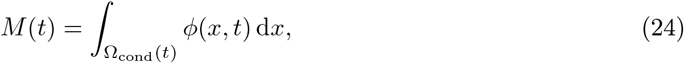

and the corresponding effective number of condensed chains is

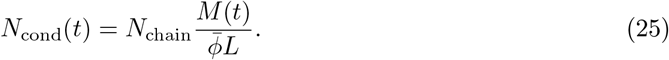

Equations (13) and (14) are solved using Fourier pseudo-spectral discretization and semi-implicit time stepping. We set *κ*_*i*_ = 2.5 × 10^−3^ for the four non-solvent fields. The periodic domain is discretized using *N* = 151 grid points, with spacing Δ*x* = *L/N* = 7.68 × 10^−3^ and a time step Δ*t* = 3.5 × 10^−6^. The model parameters and their origin are listed in Table 1.

#### Phase Classification

Each (*N*_k_, *N*_p_) was run for 90 *µ*s from a condensate slab with the enzymes at equilibrium partitioning. Over the last 40% of a run, *M* (*t*) (Eq. (24)) classifies it as dissolved (*M* < 10^−4^), oscillating (amplitude > 2 × 10^−3^, sustained to within a factor of two between the second and last quarter, period between 1 and 20 *µ*s), or stable condensate otherwise; the period is the first autocorrelation maximum of *M* . The cycle does not depend on the details of the initial condensate: at the six MD setups where the model oscillates, varying the initial condensate size, phosphorylation level, number of condensates, or enzyme distribution leads to the same limit cycle (Supplementary Fig. 29).

## Data availability

Data and code for analysis and plots, together with example trajectories and Supplementary Movies, are available as a Zenodo repository^82^ (https://doi.org/10.5281/zenodo.24088578). Additional trajectories are available at https://doi.org/10.5281/zenodo.24103647. Complete trajectories are available on request.

## Code availability

The code used for the simulations and the HOOMD-blue plugin with the Ashbaugh-Hatch pair potential for the non-bonded interactions are available on GitHub (https://github.com/ezippo/hoomd3_phosphorylation/releases/tag/v3.1_kinase_phosphatase, https://github.com/ezippo/ashbaugh_plugin). The code used for phase-field simulations is available at https://github.com/leqiao/tdp43-condensate-oscillations-phase-field.

## Acknowledgements

This project was funded by the Deutsche Forschungsgemeinschaft (DFG, German Research Foundation) - Project numbers 233630050, 465145163, and 464588647 (SFB TRR 146, project A7; SFB 1552, project C2; SFB 1551). L.S.S. thank M^3^ODEL and ReALity (Resilience, Adaptation and Longevity) and Forschungsinitiative des Landes Rheinland-Pfalz for support. The authors gratefully acknowledge the computing time provided to them at the NHR Center NHR@SW at Johannes Gutenberg University Mainz (project number 29022). This is funded by the Federal Ministry of Education and Research, and the state governments participating on the basis of the resolutions of the GWK for national high performance computing at universities (www.nhr-verein.de/unsere-partner). We thank Ellen Meyberg and Emre Pekbilir for insightful discussions.

## Author contributions

E.Z. ran the MD simulations, analyzed data, interpreted results, wrote the manuscript. L.Q. designed and ran phase field simulations, wrote the manuscript. D.D. provided important intellectual knowledge and assisted in designing the study, reviewed the manuscript. T.S. provided important intellectual knowledge and assisted in designing the study, reviewed the manuscript. F. S. supervised the study, designed phase field simulations, reviewed manuscript. L.S.S. conceived and supervised the study, interpreted results, wrote and reviewed manuscript.

## Competing interests

The authors declare no competing interests.

## Notes

### Competing Interest Statement

The authors have declared no competing interest.

### Summary of Updates

Corresponding author and minor mistakes and inconsistencies in the text.

## References

1. Kholodenko, B. N. Cell-signalling dynamics in time and space. Nature reviews Molecular cell biology 7, 165–176 (2006).

2. Hyman, A. A., Weber, C. A. & Jülicher, F. Liquid-liquid phase separation in biology. Annual review of cell and developmental biology 30, 39–58 (2014).

3. Shin, Y. & Brangwynne, C. P. Liquid phase condensation in cell physiology and disease. Science 357, eaaf4382 (2017).

4. Kota, D., Prasad, R. & Zhou, H.-X. Adenosine Triphosphate Mediates Phase Separation of Disordered Basic Proteins by Bridging Intermolecular Interaction Networks. Journal of the American Chemical Society 146. PMID: 38174879, 1326–1336 (2024).

5. Chen, M. W. et al. Biomolecular condensates can function as inherent catalysts. Molecular Cell (2026).

6. Banani, S. F., Lee, H. O., Hyman, A. A. & Rosen, M. K. Biomolecular condensates: organizers of cellular biochemistry. Nature reviews Molecular cell biology 18, 285–298 (2017).

7. Boeynaems, S. et al. Protein Phase Separation: A New Phase in Cell Biology. Trends in Cell Biology 28, 420–435 (2018).

8. Dillenburg, R. F. et al. A Phosphorylation-Induced Micellization Switch in the Low-Complexity Domain of TDP-43. Advanced Science n/a, e76625.

9. Weber, C. A., Zwicker, D., Jülicher, F. & Lee, C. F. Physics of active emulsions. Reports on Progress in Physics 82, 064601 (2019).

10. Bauermann, J., Weber, C. A. & Jülicher, F. Energy and Matter Supply for Active Droplets. Annalen der Physik 534, 2200132 (2022).

11. De, S. & Klajn, R. Dissipative self-assembly driven by the consumption of chemical fuels. Advanced Materials 30, 1706750 (2018).

12. Wanzke, C., Jussupow, A., Kohler, F., Dietz, H., Kaila, V. R. & Boekhoven, J. Dynamic vesicles formed by dissipative self-assembly. ChemSystemsChem 2, e1900044 (2020).

13. Dai, K. et al. Regulating Chemically Fueled Peptide Assemblies by Molecular Design. Journal of the American Chemical Society 142, 14142–14149 (2020).

14. Donau, C. et al. Active coacervate droplets as a model for membraneless organelles and protocells. Nature communications 11, 5167 (2020).

15. Abbas, M., Lipiński, W. P., Wang, J. & Spruijt, E. Peptide-based coacervates as biomimetic protocells. Chemical Society Reviews 50, 3690–3705 (2021).

16. Saleh, O. A., Wilken, S., Squires, T. M. & Liedl, T. Vacuole dynamics and popping-based motility in liquid droplets of DNA. Nature communications 14, 3574 (2023).

17. Zwicker, D., Decker, M., Jaensch, S., Hyman, A. A. & Jülicher, F. Centrosomes are autocatalytic droplets of pericentriolar material organized by centrioles. Proceedings of the National Academy of Sciences 111, E2636–E2645 (2014).

18. Nakashima, K. K., van Haren, M. H. I., André, A. A. M., Robu, I. & Spruijt, E. Active Coacervate Droplets Are Protocells That Grow and Resist Ostwald Ripening. Nat. Commun. 12, 3819 (2021).

19. Berthin, R., Fries, J. D., Jardat, M., Dahirel, V. & Illien, P. Microscopic and stochastic simulations of chemically active droplets. Phys. Rev. E 111, L023403 (2 2025).

20. Hertäg, K. et al. Active field theory approach to explain size control of transcriptional condensates. Macromolecular Bioscience 26, e70268 (2026).

21. Qiao, L., Gispert, P., Stelzl, L. S. & Schmid, F. Active transport as a mechanism of microphase selection in biomolecular condensates. Phys. Rev. Res. 8, 033345 (2026).

22. Lavagna, E., Delfino, F., Koniukov, G., Paloni, M., Ciandrini, L. & Barducci, A. Uncovering the thermodynamic principles of enzymatic regulation in biomolecular condensates with reactive simulations. bioRxiv (2025).

23. Meyberg, E., Robinson, J. F. & Speck, T. Impact of currents on non-equilibrium coexistence in chemically driven mixtures. arXiv preprint arXiv:2603.12481 (2026).

24. Dullweber, T., Belousov, R., Autorino, C., Petridou, N. I. & Erzberger, A. Shape Switching and Tunable Oscillations of Adaptive Droplets. Phys. Rev. Lett. 135, 128404 (2025).

25. Kovach, S. & GrandPre, T. Memory-Driven Self-Propulsion and Flocking of Chemically Active Droplets. arXiv preprint arXiv:2607.14451 (2026).

26. Datt, C., Bauermann, J., Budanur, N. B. & Jülicher, F. Fluid Flow and Spatiotemporal Chaos in Chemically Active Emulsions. Phys. Rev. Lett. 136, 254001 (25 2026).

27. Haugerud, I. S., Vuijk, H. D., Boekhoven, J. & Weber, C. A. Excitability and Oscillations of Active Droplets. Phys. Rev. Lett. 136, 218401 (2026).

28. Deng, J. & Walther, A. Programmable ATP-Fueled DNA Coacervates by Transient Liquid-Liquid Phase Separation. Chem 6, 3329–3343 (2020).

29. Sastre, J. et al. Size Control and Oscillations of Active Droplets in Synthetic Cells. Nat. Commun. 16, 2003 (2025).

30. Qian, H. Phosphorylation Energy Hypothesis: Open Chemical Systems and Their Biological Functions. Annual Review of Physical Chemistry 58. PMID: 17059360, 113–142 (2007).

31. Poprawa, S. M., Stasi, M., Kriebisch, B. A. K., Wenisch, M., Sastre, J. & Boekhoven, J. Active Droplets through Enzyme-Free, Dynamic Phosphorylation. Nat. Commun. 15, 4204 (2024).

32. Perdikari, T. M., Jovic, N., Dignon, G. L., Kim, Y. C., Fawzi, N. L. & Mittal, J. A predictive coarse-grained model for position-specific effects of post-translational modifications. Biophys. J. 120, 1187–1197 (2021).

33. Hernandez-Armendariz, A. et al. A liquid-like coat mediates chromosome clustering during mitotic exit. Molecular Cell 84, 3254–3270 (2024).

34. Mosna, S. & Dormann, D. TDP-43 Phosphorylation: Pathological Modification or Protective Factor Antagonizing TDP-43 Aggregation in Neurodegenerative Diseases? Bioessays 48, e70084 (2026).

35. Gruijs da Silva, L. A. et al. Disease-linked TDP-43 hyperphosphorylation suppresses TDP-43 condensation and aggregation. EMBO J. 41, e108443.

36. Ambadipudi, S., Biernat, J., Riedel, D., Mandelkow, E. & Zweckstetter, M. Liquid-liquid phase separation of the microtubule-binding repeats of the Alzheimer-related protein Tau. Nature Communications 8, 275 (2017).

37. Wegmann, S. et al. Tau protein liquid–liquid phase separation can initiate tau aggregation. The EMBO journal 37, EMBJ201798049 (2018).

38. Buratti, E. TDP-43 post-translational modifications in health and disease. Expert Opinion on Therapeutic Targets 22. PMID: 29431050, 279–293 (2018).

39. Yamazaki, H., Takagi, M., Kosako, H., Hirano, T. & Yoshimura, S. H. Cell Cycle-Specific Phase Separation Regulated by Protein Charge Blockiness. Nat. Cell Biol. 24, 625–632 (2022).

40. He, S. et al. Kinase KEY1 Controls Pyrenoid Condensate Size throughout the Cell Cycle by Disrupting Phase Separation Interactions. Nat. Cell Biol., 1–14 (2026).

41. Novák, B. & Tyson, J. J. Design Principles of Biochemical Oscillators. Nat. Rev. Mol. Cell Biol. 9, 981–991 (2008).

42. Goldbeter, A. & Koshland, D. E. An Amplified Sensitivity Arising from Covalent Modification in Biological Systems. Proc. Natl. Acad. Sci. U. S. A. 78, 6840–6844 (1981).

43. Tung, H.-R. Precluding oscillations in Michaelis–Menten approximations of dual-site phosphorylation systems. Mathematical Biosciences 306, 56–59 (2018).

44. Nakajima, M. et al. Reconstitution of Circadian Oscillation of Cyanobacterial KaiC Phosphorylation in Vitro. Science 308, 414–415 (2005).

45. Kimchi, O. et al. Self-Assembly–Based Posttranslational Protein Oscillators. Sci. Adv. 6, eabc1939 (2020).

46. Loose, M., Fischer-Friedrich, E., Ries, J., Kruse, K. & Schwille, P. Spatial Regulators for Bacterial Cell Division Self-Organize into Surface Waves in Vitro. Science 320, 789–792 (2008).

47. Liachko, N. F. et al. The phosphatase calcineurin regulates pathological TDP-43 phosphorylation. Acta neuropathologica 132, 545–561 (2016).

48. Wang, J. et al. A molecular grammar governing the driving forces for phase separation of prion-like RNA binding proteins. Cell 174, 688–699 (2018).

49. Cao, F., von Bülow, S., Tesei, G. & Lindorff-Larsen, K. A coarse-grained model for disordered and multi-domain proteins. Protein Science 33, e5172 (2024).

50. Regy, R. M., Thompson, J., Kim, Y. C. & Mittal, J. Improved coarse-grained model for studying sequence dependent phase separation of disordered proteins. Protein Science 30, 1371–1379 (2021).

51. Albaugh, A. & Gingrich, T. R. Simulating a chemically fueled molecular motor with nonequilibrium molecular dynamics. Nature communications 13, 2204 (2022).

52. Zippo, E., Dormann, D., Speck, T. & Stelzl, L. S. Molecular simulations of enzymatic phosphorylation of disordered proteins and their condensates. en. Nat Commun 16, 4649 (2025).

53. Weng, S.-L., Rekhi, S., Kim, Y. C., Palmer, J. C. & Mittal, J. Molecular origins of pH gradients in charge-regulated biomolecular condensates. The Journal of Chemical Physics 165, 114107 (2026).

54. Das, R. K. & Pappu, R. V. Conformations of intrinsically disordered proteins are influenced by linear sequence distributions of oppositely charged residues. Proceedings of the National Academy of Sciences 110, 13392–13397 (2013).

55. Firman, T. & Ghosh, K. Sequence charge decoration dictates coil-globule transition in intrinsically disordered proteins. The Journal of chemical physics 148 (2018).

56. Zwicker, D., Hyman, A. A. & Jülicher, F. Suppression of Ostwald ripening in active emulsions. Phys. Rev. E 92, 012317 (1 2015).

57. Bauermann, J., Bartolucci, G. & Kolchinsky, A. Spatiotemporal Organization of Chemical Oscillators via Phase Separation. Phys. Rev. Lett. 136, 178001 (17 2026).

58. Speck, T. Modeling of biomolecular machines in non-equilibrium steady states. The Journal of Chemical Physics 155, 230901 (2021).

59. Dignon, G. L., Zheng, W., Kim, Y. C., Best, R. B. & Mittal, J. Sequence determinants of protein phase behavior from a coarse-grained model. PLoS Comput. Biol. 14, e1005941 (2018).

60. Tesei, G. & Lindorff-Larsen, K. Improved predictions of phase behaviour of intrinsically disordered proteins by tuning the interaction range [version 2; peer review: 2 approved]. Open Research Europe 2 (2023).

61. De Los Rios, P. & Barducci, A. Hsp70 chaperones are non-equilibrium machines that achieve ultra-affinity by energy consumption. eLife 3 (ed Chakraborty, A. K.) e02218 (2014).

62. Liu, B., Xue, M., Qiu, Y., Konovalov, K. A., O’Connor, M. S. & Huang, X. GraphVAMPnets for uncovering slow collective variables of self-assembly dynamics. The Journal of Chemical Physics 159, 094901 (2023).

63. Zhu, Y., Degiacomi, M. T., Mey, A. S. & Singh, S. Tyrosine kinases sample unique activation ensembles. bioRxiv, 2026–06 (2026).

64. Wang, J. T. et al. Regulation of RNA granule dynamics by phosphorylation of serine-rich, intrinsically disordered proteins in C. elegans. eLife 3 (ed Nilsen, T.) e04591 (2014).

65. Wippich, F., Bodenmiller, B., Trajkovska, M. G., Wanka, S., Aebersold, R. & Pelkmans, L. Dual Specificity Kinase DYRK3 Couples Stress Granule Condensation/Dissolution to mTORC1 Signaling. Cell 152, 791–805 (2013).

66. Wurtz, J. D. & Lee, C. F. Stress granule formation via ATP depletion-triggered phase separation. New Journal of Physics 20, 045008 (2018).

67. Liu, J.-L., Hebert, M. D., Ye, Y., Templeton, D. J.Kung, H.-J. & Matera, A. G. Cell cycle-dependent localization of the CDK2-cyclin E complex in Cajal (coiled) bodies. Journal of Cell Science 113, 1543–1552 (2000).

68. Hnisz, D., Shrinivas, K., Young, R. A., Chakraborty, A. K. & Sharp, P. A. A Phase Separation Model for Transcriptional Control. Cell 169, 13–23 (2017).

69. Anderson, J. A., Glaser, J. & Glotzer, S. C. HOOMD-blue: A Python package for high-performance molecular dynamics and hard particle Monte Carlo simulations. Computational Materials Science 173, 109363 (2020).

70. Rauh, A. S., Hedemark, G. S., Tesei, G. & Lindorff-Larsen, K. A coarse-grained model for simulations of phosphorylated disordered proteins. Biophysical Journal 125, 396–405 (2026).

71. Gemeinhardt, T. M. et al. A disordered linker in the Polycomb protein Polyhomeotic tunes phase separation and oligomerization. Molecular cell 85, 2128–2146 (2025).

72. Gebel, J. et al. p63 uses a switch-like mechanism to set the threshold for induction of apoptosis. Nature Chemical Biology 16, 1078–1086 (2020).

73. Hendus-Altenburger, R. et al. Molecular basis for the binding and selective dephosphorylation of Na+/H+ exchanger 1 by calcineurin. Nature communications 10, 3489 (2019).

74. Metropolis, N., Rosenbluth, A. W., Rosenbluth, M. N., Teller, A. H. & Teller, E. Equation of state calculations by fast computing machines. The journal of chemical physics 21, 1087–1092 (1953).

75. Buchete, N.-V. & Hummer, G. Coarse Master Equations for Peptide Folding Dynamics. The Journal of Physical Chemistry B 112. PMID: 18232681, 6057–6069 (2008).

76. Trendelkamp-Schroer, B., Wu, H., Paul, F. & Noé, F. Estimation and uncertainty of reversible Markov models. The Journal of Chemical Physics 143, 174101 (2015).

77. Hoffmann, M. et al. Deeptime: a Python library for machine learning dynamical models from time series data. Machine Learning: Science and Technology (2021).

78. Prinz, J.-H. et al. Markov models of molecular kinetics: Generation and validation. The Journal of Chemical Physics 134, 174105 (2011).

79. Wu, H. & Noé, F. Variational approach for learning Markov processes from time series data. Journal of Nonlinear Science 30, 23–66 (2020).

80. Ramasubramani, V., Dice, B. D., Harper, E. S., Spellings, M. P., Anderson, J. A. & Glotzer, S. C. freud: A Software Suite for High Throughput Analysis of Particle Simulation Data. Computer Physics Communications 254, 107275 (2020).

81. Ensign, D. L. & Pande, V. S. Bayesian Single-Exponential Kinetics in Single-Molecule Experiments and Simulations. The Journal of Physical Chemistry B 113, 12410–12423 (2009).

82. Zippo, E. Supporting Data: Molecular simulations of enzymatic phosphorylation of disordered proteins and their condensates 2024.

