## Supplementary Information for "Kinase–phosphatase cycles drive self-sustained oscillations of protein condensates"

### Supplementary Text

##### Thermodynamic consistency of molecular dynamics simulations of kinase-phosphatase cycles

To build a complete phosphorylation-dephosphorylation cycle with both kinase and phosphatase, we verified first that simulations with single enzyme and TDP-43 are thermodynamically consistent. We used the CALVADOS3 force field[1,2], with an elastic network employed to fix the secondary structure of the enzymes, which allows the partially flexible structures to undergo breathing motions around their mean conformation. In this case, we introduced an additional Metropolis move to swap Ser and pSer without driving if the distance with the enzyme active site is greater than half box size (15 nm). In this way we mimic the exchange of substrate and product with an external chemical bath at fixed concentrations. Distances were checked every 200 MD steps, as for the phosphorylation/dephosphorylation moves. In order to optimize the estimation of the MSM, we chose the most reactive serines as phosphosite (Ser395 for the simulations with calcineurin, Ser403 for the ones with CK1$\delta$). We run 5 simulation replicas for $20 \times 10^8$ MD steps (20 $\mu$s in simulation time) and dumped snapshots every 5000 MD steps. For both reaction cycles with kinase and with phosphatase, we discretized the trajectories into 4 Markov states (substrate unbound, substrate bound to enzyme, product bound to enzyme, product unbound) and computed the heat dissipated per cycle from the ratio of the product of transition rates in the forward and backward directions[3] (Methods).

The thermodynamic consistency test holds in a reaction cycle with TDP-43 LCD and calcineurin phosphatase (Supplementary Fig. 1,2,3). We performed simulations at different drivings, namely $\Delta\mu = -2.5, 0, +2.5, +5$ kJ mol$^{-1}$. The rate model for the reactive cycle shows how detailed balance is observed for all steps except for the exchange between states 2 and 3, as the chemical driving is applied at this step. The chemically-driven exchange between states 2 and 3 however obeys local-detailed balance (Supplementary Fig. 1, Methods). As an additional check, we repeated the test at $\Delta\mu = +2.5$ kJ mol$^{-1}$ using VAMPnet[4] to recognize the bound states in an automated unsupervised fashion (Supplementary Fig. 1,2,3). We followed the same architecture of our previous work[3], re-training the network on simulations with calcineurin and TDP-43.

The thermodynamic consistency test holds also in a reaction cycle with CK1$\delta$ and 5 times phosphorylated TDP-43 LCD (5p TDP-43) as substrate (Supplementary Fig. 4,5,6), as we had previously shown for a cycle with wild type TDP-43 LCD and with a rigid representation of the kinase structure[3].

##### Comparison of CALVADOS3 and HPS-Urry coarse-grained simulations

The CALVADOS3[1,2] and HPS-Urry[5–7] simulation models enable accurate simulations of multi-domain proteins. The HPS-Urry simulations show stronger propensity to phase separate at simulation unit room temperature. To compare the two simulation models we thus ran simulations with the CALVADOS3 model at lower temperature. First we simulated each enzyme in absence of phosphorylation. For calcineurin, we find that the enzyme prefers the condensate interface over the condensate interior in simulations with both CALVADOS3 and HPS-Urry models. Note that the HPS-Urry profile is not symmetric, which is due to the finite simulation time. In CALVADOS3, there is more calcineurin in the dilute solution than in simulations with HPS-Urry (Supplementary Fig. 11). When pSer are present, calcineurin is mostly interfacial or in the dilute phase in simulations with CALVADOS3. In simulation with HPS-Urry, calcineurin is at the interface of the condensate with the dilute phase, but also enters the condensate core, as in the simulations with unphosphorylated TDP-43 LCD condensates (Supplementary Fig. 11).

Differences in the co-condensation profiles can be in part due to an imperfect matching of temperatures between the models, but may also reflect differences in the intrinsic interaction propensities of different residues considering the differences in the parameterization of the two simulation models, but overall the two simulation models provided a consistent picture, with enzymes preferentially interacting with the condensate interface for unphosphorylated TDP-43 LCD, new interactions upon phosphorylation and due to the folded domains of full-length TDP-43.

##### Fit of density profiles

We used a hyperbolic tangent fit of the residue density profiles to estimate both the residue densities in dense and dilute phase from Fig. 3c, and the interface width of the condensate in simulations at $N_{\mathrm{k}} = 2$, $N_{\mathrm{p}} = 24$ (Fig. 3g, Methods). We fitted the residue densities at different distances from the condensate center of mass $\rho(r)$ with the function:

$$\rho_{\mathrm{th}}(x) = a_0 - a_1 \tanh\left(a_2(x - a_3)\right). \tag{1}$$

We defined the densities of dense and dilute phase as $\rho_{\mathrm{dense}} = \lim_{x\to-\infty} \rho_{\mathrm{th}}(x) = a_0 + a_1$ and $\rho_{\mathrm{dilute}} = \lim_{x\to+\infty} \rho_{\mathrm{th}}(x) = a_0 - a_1$, while $w = 1/a_2$ is the width and $R_{\mathrm{cond}} = a_3$ is the condensate radius.

##### Independence of the limit cycle from the initial conditions

The state diagram of Fig. 4e is computed from a single initial condition, a condensate slab with the enzymes at equilibrium partitioning. To test whether the limit cycle depends on it, each of the six MD setups where the model oscillates was also started from 15 other initial conditions: droplets of 0.25–1 times the coexistence size, pre-phosphorylated to $p = 0.2$ or 0.4, with neutrally partitioned enzymes ($\phi_E \propto 1 - \phi$) or with the kinase confined to the dilute phase, two or three droplets of unequal size, and restarts from the maximum and minimum of the cycle. All runs converge to the same limit cycle as the slab start (Supplementary Fig. 29): the period is identical, and the mean and amplitude of the phosphorylation agree to within 0.2%.

### Supplementary Figures

#### Thermodynamic consistency simulations of driven dephosphorylation of TDP-43

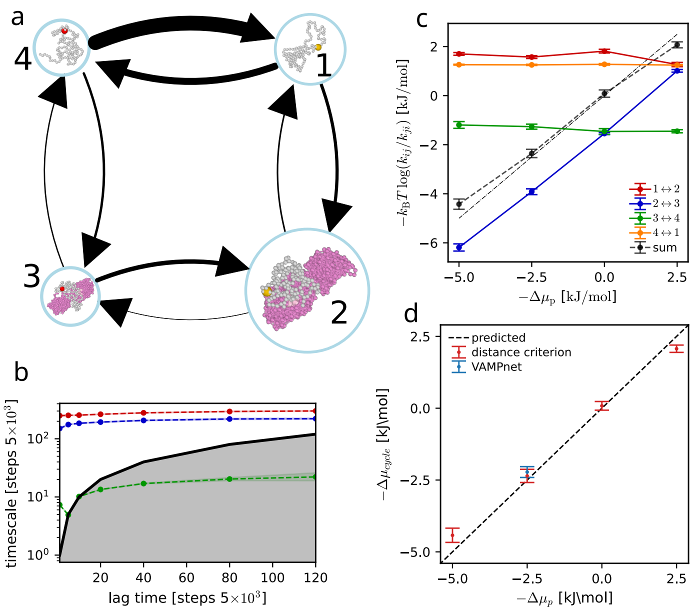


Supplementary Figure 1: (**a**) Sketch of the MSM for an example simulation with TDP-43 LCD (gray, pSer in red) and calcineurin (magenta) at $\Delta\mu_\mathrm{p} = +2.5$ kJ mol$^{-1}$. The size of the arrows is proportional to the absolute value of the corresponding transition rate and the size of the circular node is proportional to the stationary distribution of the corresponding state. (**b**) Implied timescales at different lag times for the 4-state MSM trajectory shown in **a**. Shaded areas represent 95% confidence intervals from 100 bootstrapped samples. (**c**) Mean values +/- SEM of the single edge contributions to $\Delta\mu_\mathrm{cycle}$, that is $k_B T \log(k_{ij}/k_{ji})$. (**d**) Mean values +/- SEM of $\Delta\mu_\mathrm{cycle}$ vs $\Delta\mu_\mathrm{p}$.

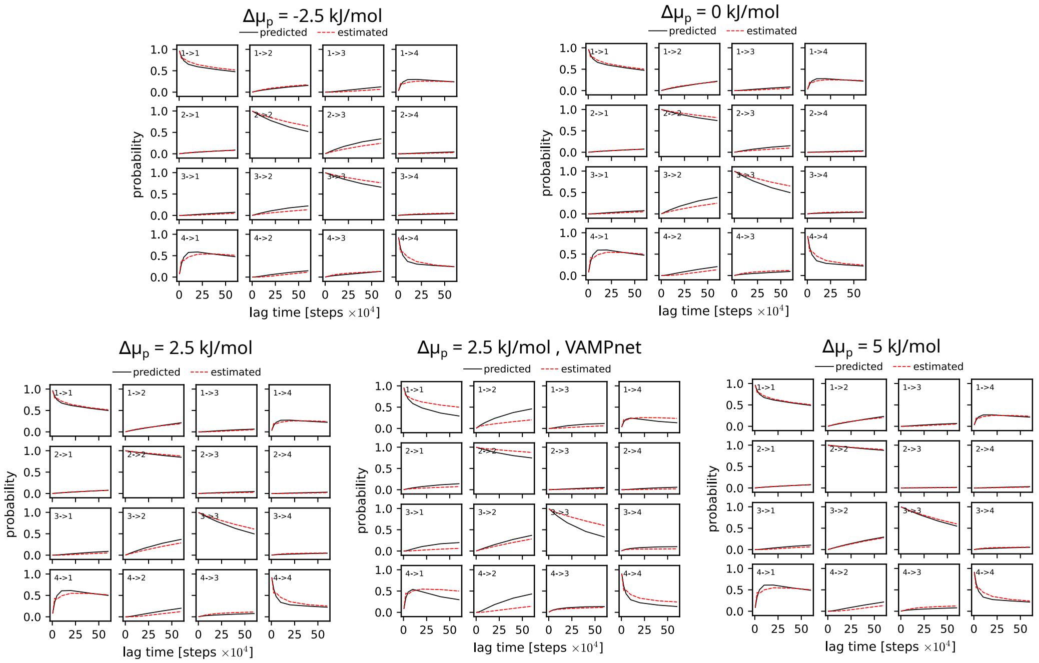


Supplementary Figure 2: CK test for 4-state cycles with TDP-43 and calcineurin at different $\Delta\mu_\mathrm{p}$. Shaded areas are 95% confidence intervals from 100 bootstrapped samples.

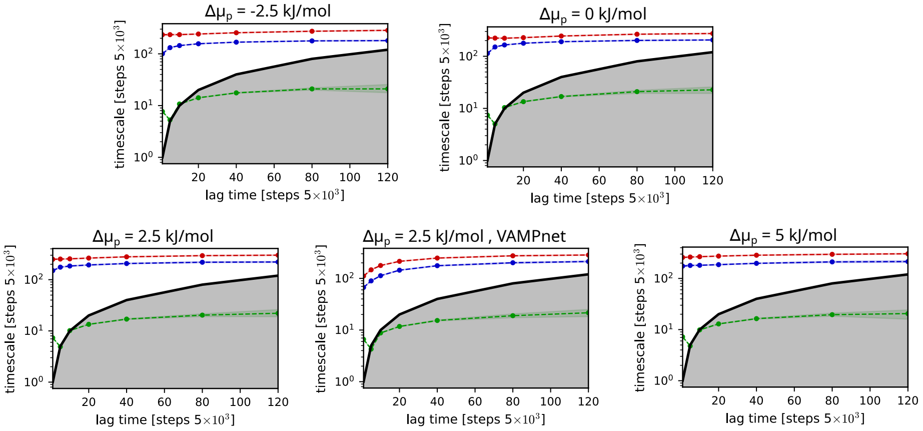


Supplementary Figure 3: Implied timescales for 4-state cycles with TDP-43 and calcineurin at different $\Delta\mu_\mathrm{p}$. Shaded areas are 95% confidence intervals from 100 bootstrapped samples.

#### Thermodynamic consistency simulations of ATP-driven phosphorylation of TDP-43

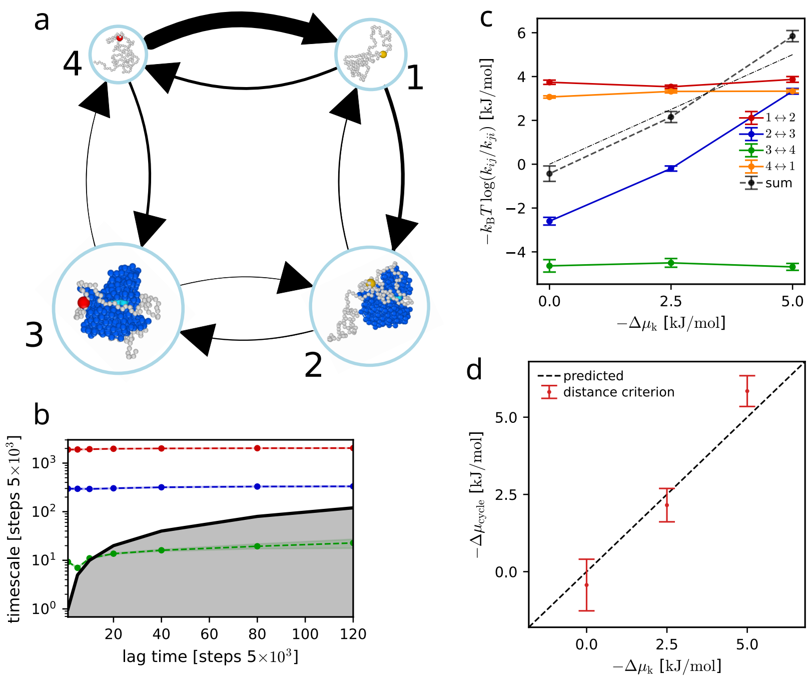


Supplementary Figure 4: (**a**) Sketch of the MSM for an example simulation with TDP-43 LCD (gray, pSer in red) and CK1$\delta$ (blue) at $\Delta\mu_\mathrm{k} = -5$ kJ mol$^{-1}$. The size of the arrows is proportional to the absolute value of the corresponding transition rate and the size of the circular node is proportional to the stationary distribution of the corresponding state. (**b**) Implied timescales at different lag times for the 4-state MSM trajectory shown in **a**. Shaded areas represent 95% confidence intervals from 100 bootstrapped samples. (**c**) Mean values +/- SEM of the single edge contributions to $\Delta\mu_\mathrm{cycle}$, that is $k_BT\log(k_{ij}/k_{ji})$. (**d**) Mean values +/- SEM of $\Delta\mu_\mathrm{cycle}$ vs $\Delta\mu_\mathrm{k}$.

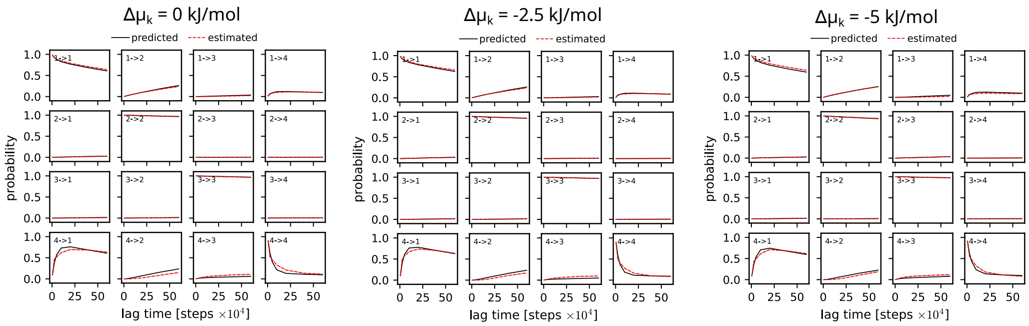


Supplementary Figure 5: CK test for 4-state cycles with TDP-43 and CK1$\delta$ at different $\Delta\mu_\mathrm{k}$. Shaded areas are 95% confidence intervals from 100 bootstrapped samples.

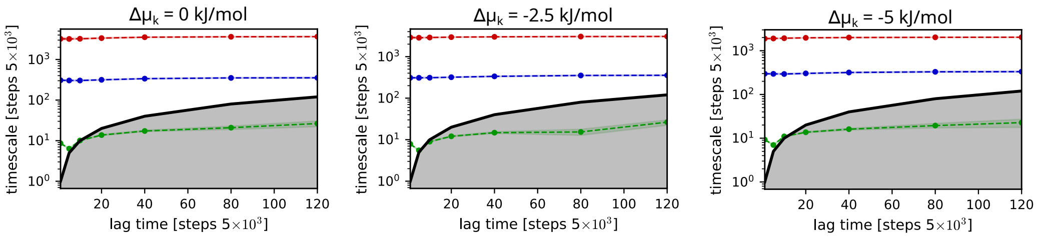


Supplementary Figure 6: Implied timescales for 4-state cycles with TDP-43 and CK1$\delta$ at different $\Delta\mu_\mathrm{k}$. Shaded areas are 95% confidence intervals from 100 bootstrapped samples.

#### Thermodynamic consistency simulations of kinase-phosphatase cycles

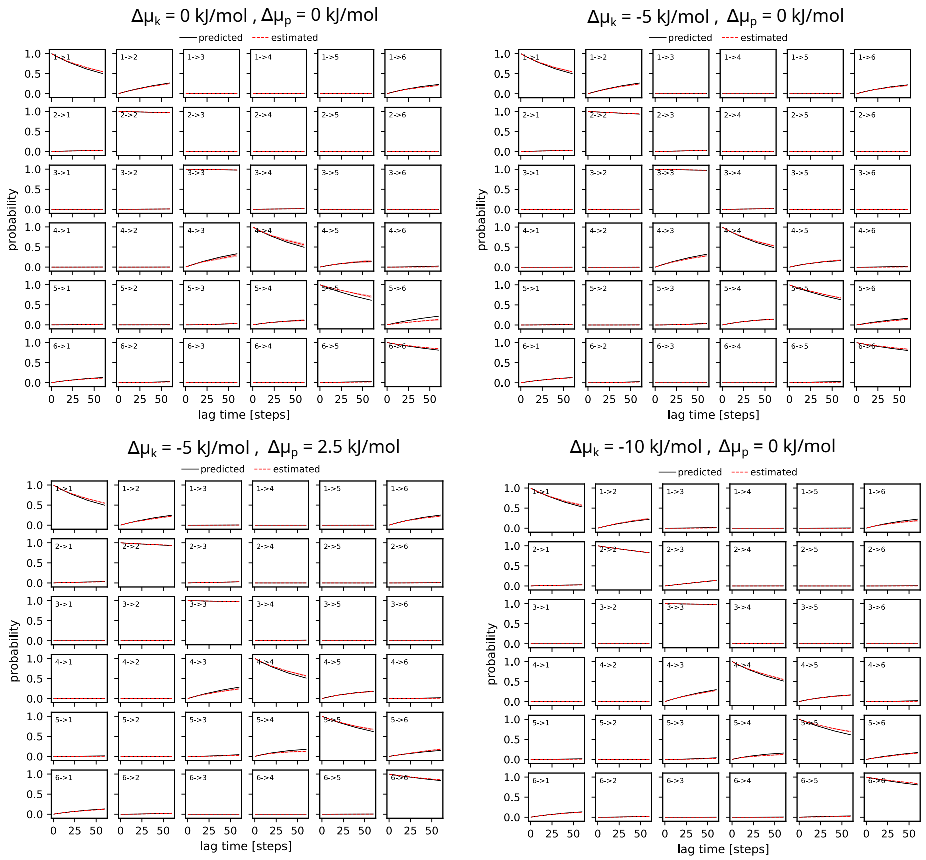


Supplementary Figure 7: CK test for 6-state cycles with TDP-43, calcineurin and CK1$\delta$ at different $\Delta\mu_\mathrm{k}$, $\Delta\mu_\mathrm{p}$. Shaded areas are 95% confidence intervals from 100 bootstrapped samples.

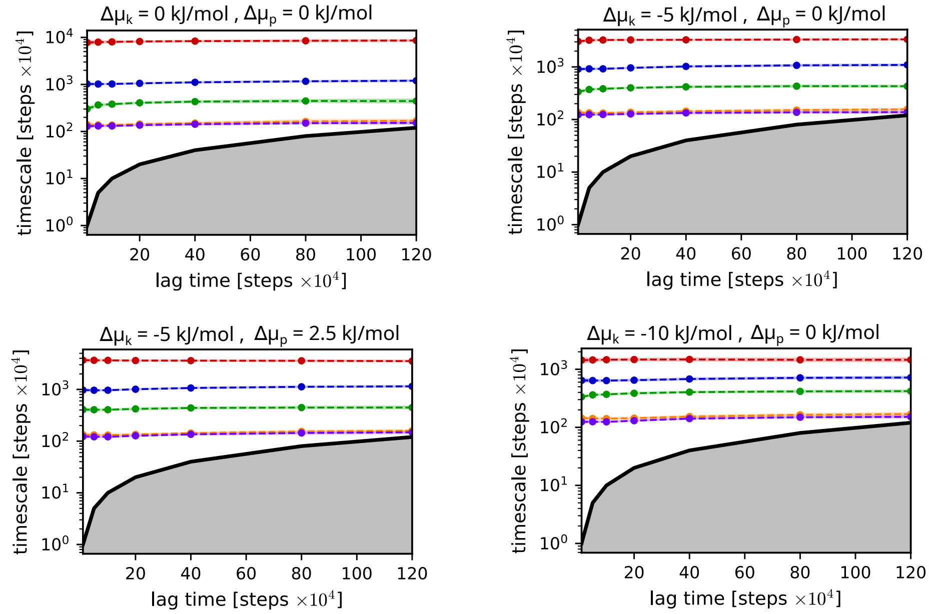


Supplementary Figure 8: Implied timescales for 6-state cycles with TDP-43, calcineurin and CK1$\delta$ at different $\Delta\mu_{\mathrm{k}}$, $\Delta\mu_{\mathrm{p}}$. Shaded areas are 95% confidence intervals from 100 bootstrapped samples.

#### Condensate and enzymes organization

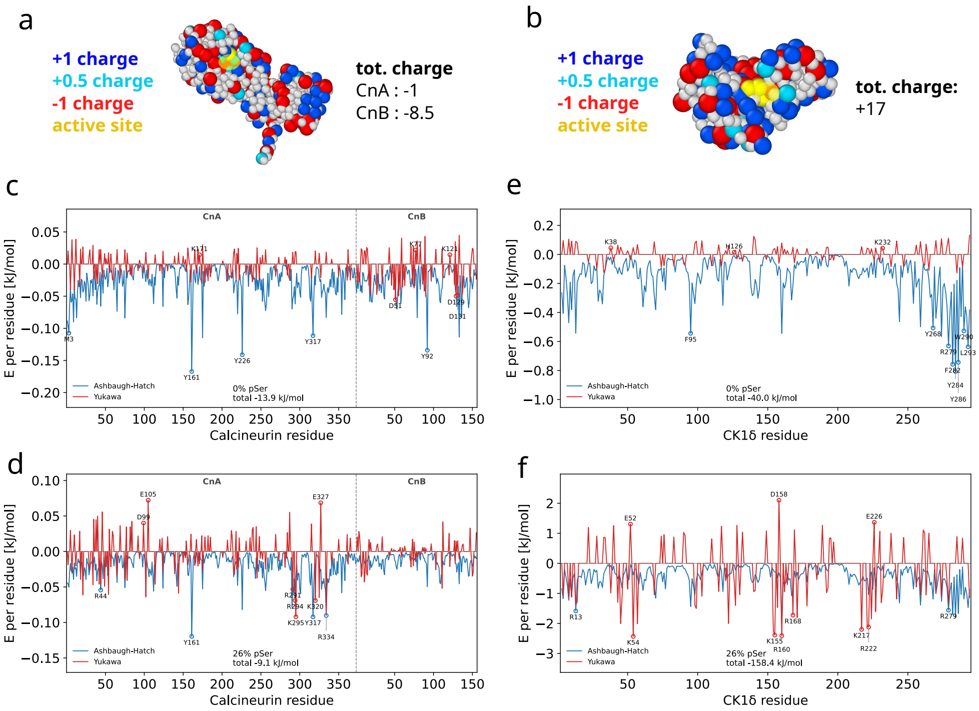


Supplementary Figure 9: (**a-b**) Residue-level coarse-grained illustration of calcineurin (a) and CK1$\delta$ (b), with colored charged residues (-1 red, +1 blue, +0.5 light blue) and active site (yellow). (**c-f**) Ashbaugh-Hatch (blue) and Yukawa (red) energy contributions for each residue of calcineurin (c,d) and CK1$\delta$ (e,f) in the interaction with TDP-43 LCD residues, from equilibrium simulations in slab geometry both without phosphorylations (c,e) and with 26% of phosphorylated Ser (d,f).

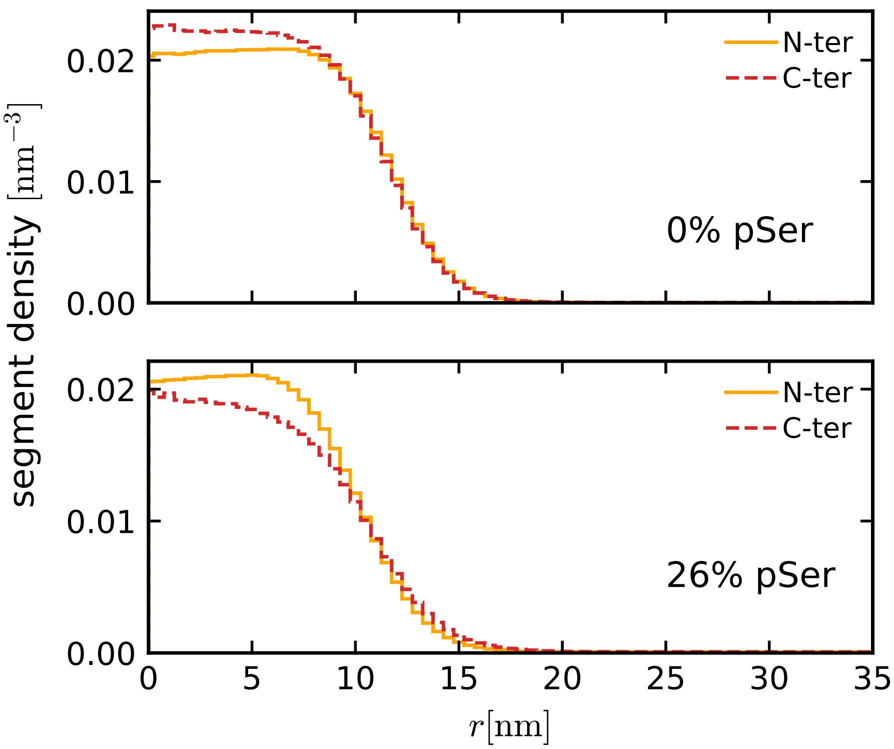


Supplementary Figure 10: N-terminal (residues 261-360, orange) and C-terminal (residues 361-414, red) segments density profiles from equilibrium simulation with 200 TDP-43 LCD wild type (top) or with 26% pSer (bottom) and for 5 calcineurin (same as in Fig. 2d in main text).

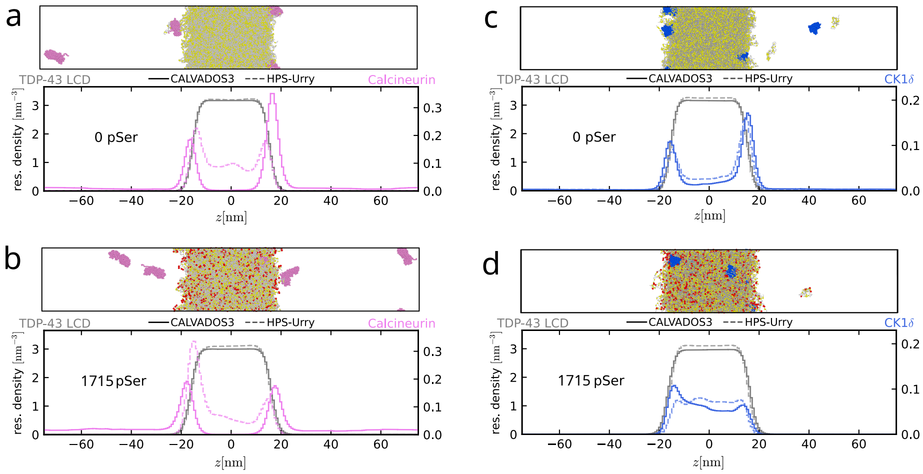


Supplementary Figure 11: Residue density profile for a condensate of 400 TDP-43 LCD (gray), either wild type (**a**,**c**) or with 1715 pSer (**b**,**d**), and for 5 enzymes, either calcineurin (**a**,**b**, magenta) or CK1$\delta$ (**c**,**d**, blue). We compare CALVADOS3 (solid lines) and HPS-Urry (dashed lines) force fields. Data from 15 $\mu$s long trajectories with pre-equilibrated condensate. On top of each plot, snapshot from the CALVADOS3 simulation. TDP-43 is gray, with yellow Ser and red pSer, calcineurin are magenta, CK1$\delta$ are blue.

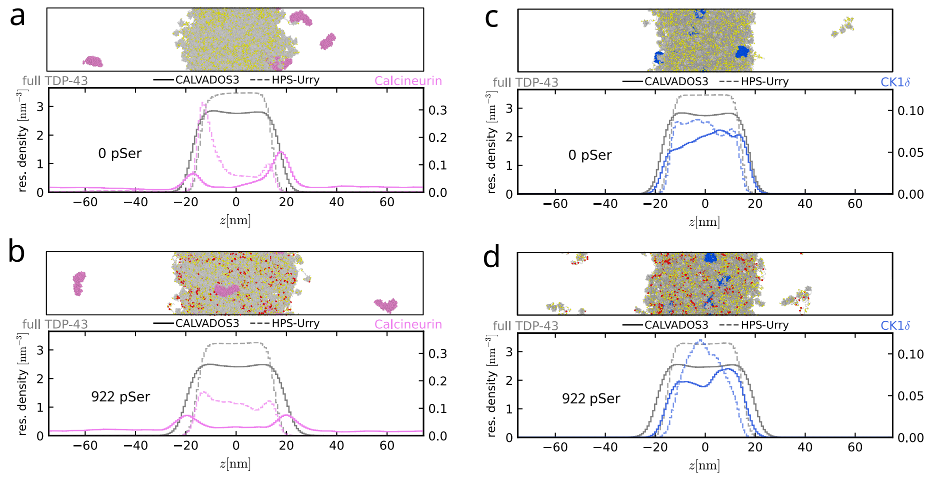


Supplementary Figure 12: Residue density profile for a condensate of 150 full-length TDP-43 (gray), either wild type (**a**,**c**) or with 922 pSer (**b**,**d**), and 5 enzymes, either calcineurin (**a**,**b**, magenta) or CK1$\delta$ (**c**,**d**, blue). We compare CALVADOS3 (solid lines) and HPS-Urry (dashed lines) force fields. Data from 15 $\mu$s long trajectories with pre-equilibrated condensate. On top of each plot, snapshot from the CALVADOS3 simulation. TDP-43 is gray, with yellow Ser and red pSer, calcineurin are magenta, CK1$\delta$ are blue.

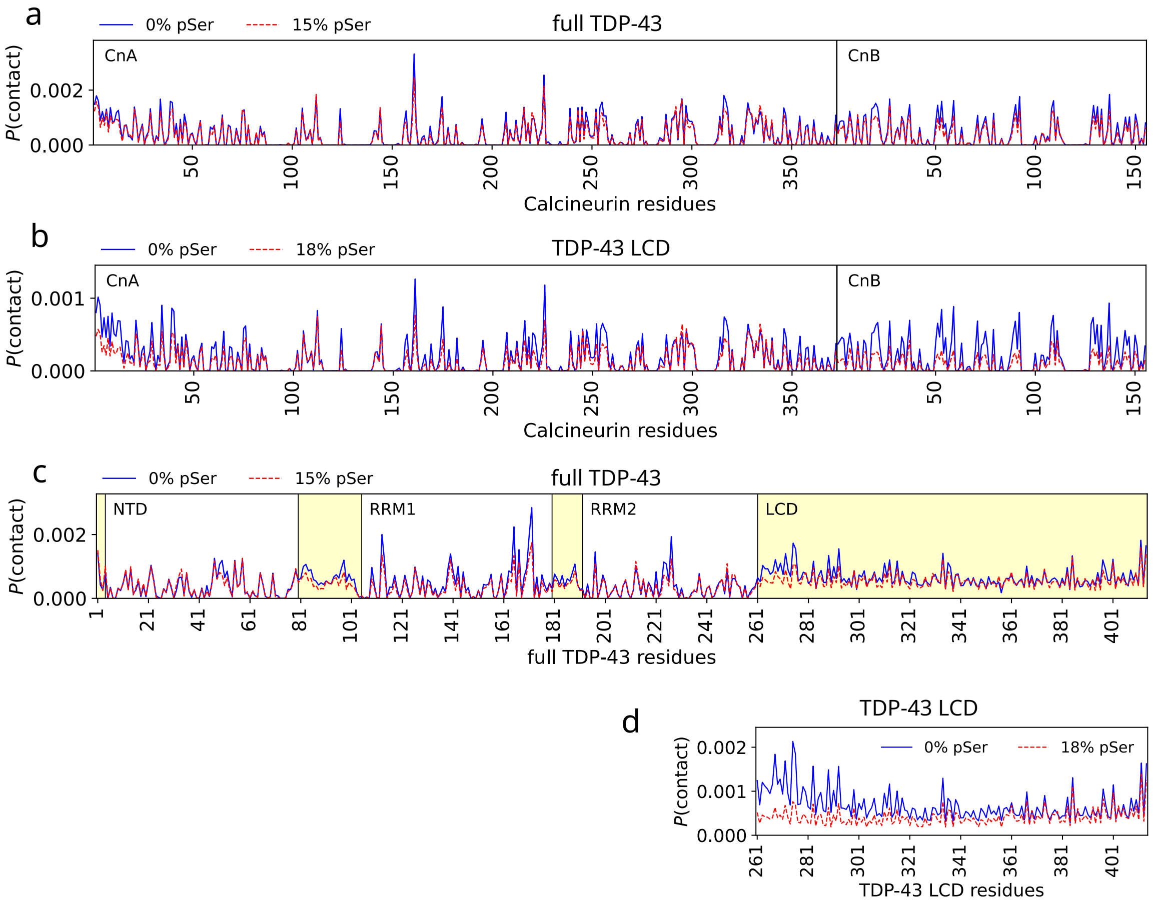


Supplementary Figure 13: (**a,b**) Probability of contact between each residue of calcineurin and full length TDP-43 (a) or TDP-43 LCD (b) chains in one snapshot. (**c,d**) Probability of contact between each residue of full length TDP-43 (c) or TDP-43 LCD (d) with calcineurin chains in one snapshot.

#### Phosphorylation patterns

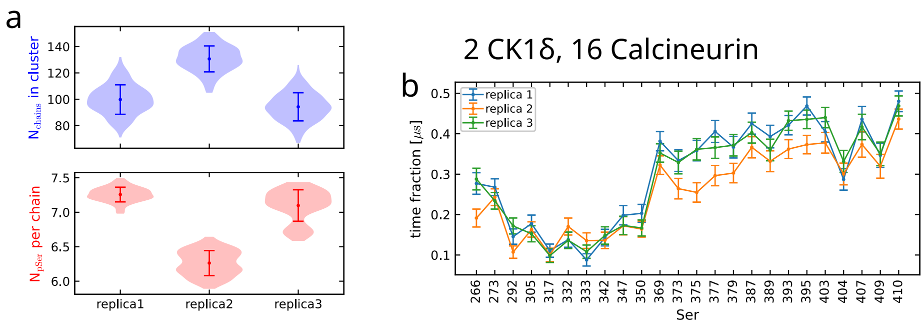


Supplementary Figure 14: (**a**) Distributions with mean and standard deviation of number of chains in biggest cluster (blue, top) and number of pSer per chain (red, bottom) for 3 replicas at $\mathrm{N_k} = 2$, $\mathrm{N_p} = 16$. (**b**) Time fraction in phosphorylated state for each Ser residue averaged from all chains in simulation for 3 different replicas. Errors computed with bootstrap method.

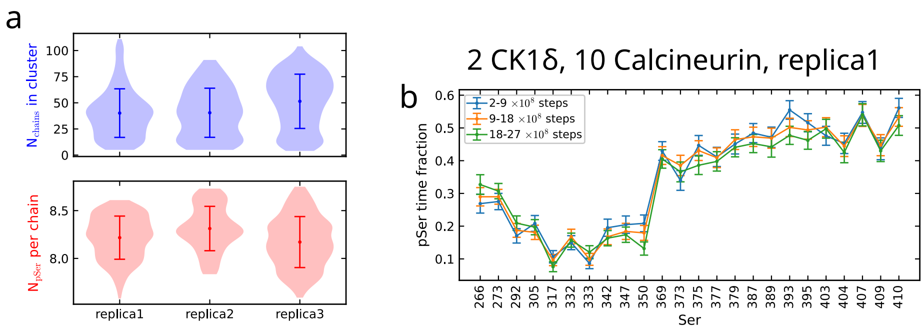


Supplementary Figure 15: (**a**) Distributions with mean and standard deviation of number of chains in biggest cluster (blue, top) and number of pSer per chain (red, bottom) for 3 replicas at $N_\mathrm{k} = 2$, $N_\mathrm{p} = 10$. (**b**) Time fraction in phosphorylated state for each Ser residue averaged from all chains in simulation for the first 3 oscillation periods of replica 1. Errors computed with bootstrap method.

#### Reactions at interface

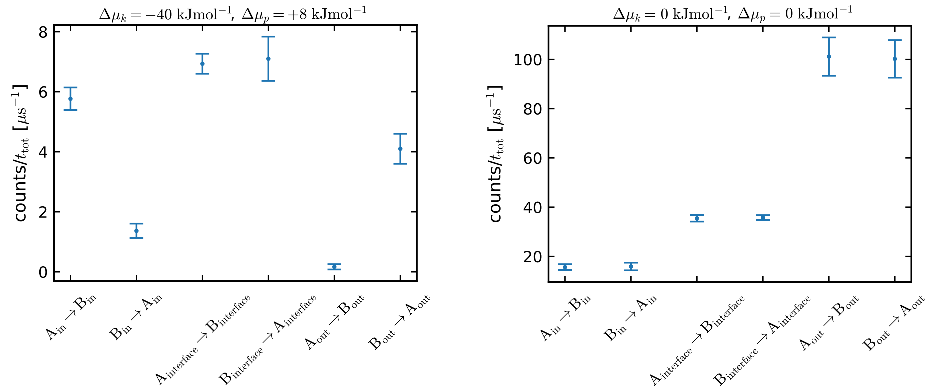


Supplementary Figure 16: Frequency of transitions between chemical states A and B in condensate core, at interface and in the dilute phase. Errors are computed from the 3 different replicas.

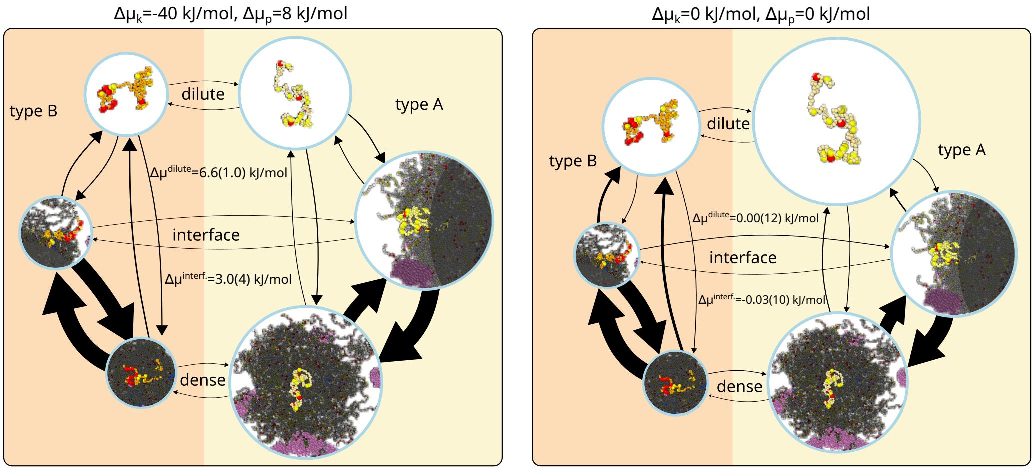


Supplementary Figure 17: MSM cycle sketches showing transitions from chains of type A (low phosphorylation degree, yellow chain) to B (high phosphorylation degree, orange chain) and between dense, dilute and interface regions, from simulations with 2 CK1$\delta$ and 24 calcineurin with (left, $\Delta\mu_{\rm k} = -40$ kJ mol$^{-1}$, $\Delta\mu_{\rm p} = +8$ kJ mol$^{-1}$) and without driving (right, $\Delta\mu_{\rm k} = 0$ kJ mol$^{-1}$, $\Delta\mu_{\rm p} = 0$ kJ mol$^{-1}$). The size of the arrows is proportional to the absolute value of the corresponding transition rate and the size of the circular node is proportional to the stationary distribution of the corresponding state.

#### Reactions in passive system

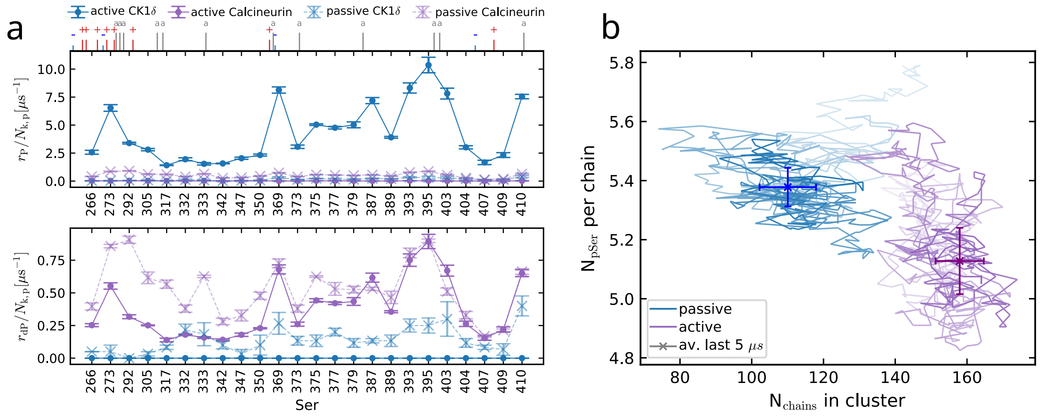


Supplementary Figure 18: (**a**) Phosphorylation (up) and dephosphorylation (bottom) rates for each Ser residue by CK1$\delta$ and calcineurin from 3 replicas at Nk = 2, Np = 24, both with and without driving. (**b**) Phase portrait with number of chains in condensate on x-axis and number of pSer per chain in y-axis, for simulations with $N_{\mathrm{k}} = 2$, $N_{\mathrm{p}} = 24$. One example trajectory at $\Delta\mu_{\mathrm{k}} = 0$ kJ mol$^{-1}$ and $\Delta\mu_{\mathrm{p}} = 0$ kJ mol$^{-1}$ (blue) and one at $\Delta\mu_{\mathrm{k}} = -40$ kJ mol$^{-1}$ and $\Delta\mu_{\mathrm{p}} = 8$ kJ mol$^{-1}$ (purple). Cross markers indicate mean and standard deviation from the last $5\times10^{8}$ MD steps.

#### Ostwald ripening

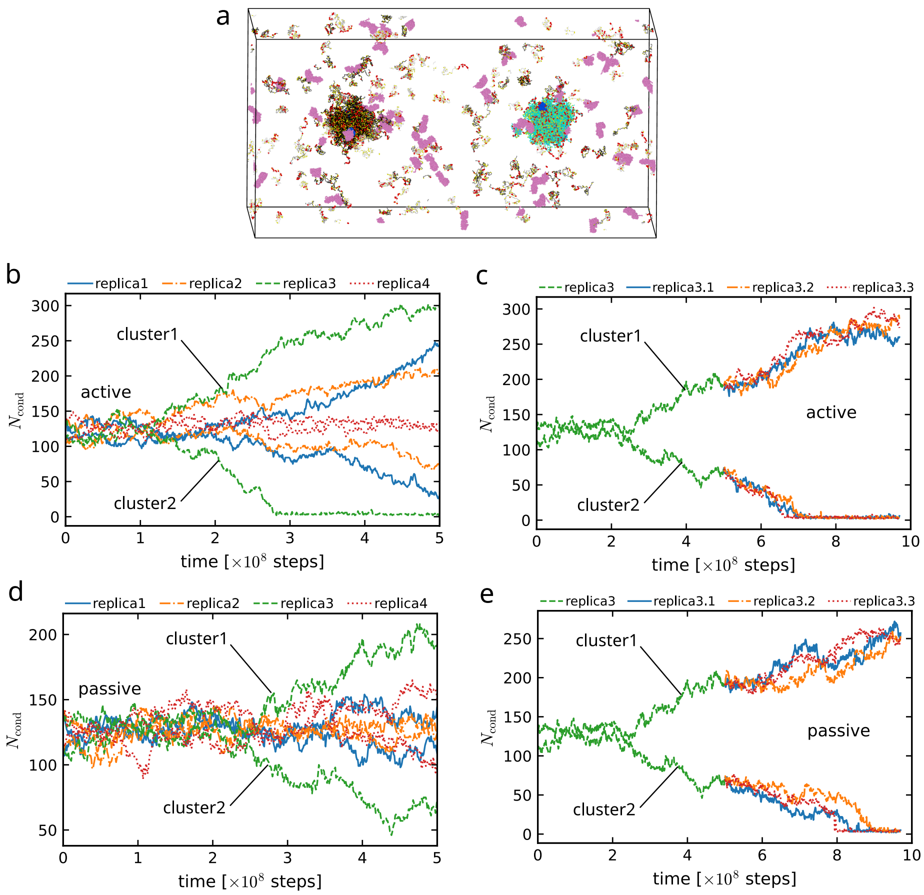


Supplementary Figure 19: (**a**) Starting configuration, merging two snapshots from simulation at $N_{\rm k} = 2, N_{\rm p} = 24$, $\Delta\mu_{\rm k} = -40$ kJ mol$^{-1}$, $\Delta\mu_{\rm p} = 8$ kJ mol$^{-1}$. Cyan and black chains represent respectively the biggest and second biggest cluster. CK1$\delta$ are in blue, calcineurin in magenta, Ser and pSer residues in yellow and red respectively. (**b-e**) $N_{\rm cond}$ in time for the two biggest clusters from 4 simulation replicas of 5$\mu$s with active reactions at $\Delta\mu_{\rm k} = -40$ kJ mol$^{-1}$, $\Delta\mu_{\rm p} = 8$ kJ mol$^{-1}$ (b) and without reactions (d). Starting from the partially equilibrated replica 3 without reactions, with a big cluster and a small cluster, we run 3 more replicas with active reactions (c) and with no reactions (e).

#### Delay from kinase binding

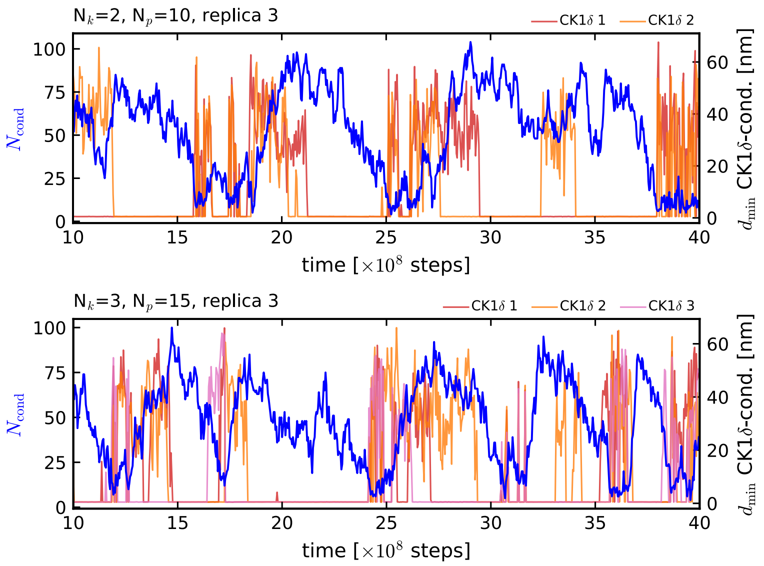


Supplementary Figure 20: Trajectories of condensate size (blue, left y-axis) and minimum distance between each CK1$\delta$ and the biggest cluster in the simulation (red, right y-axis) for $N_{\rm k} = 2$, $N_{\rm p} = 10$ and $N_{\rm k} = 3$, $N_{\rm p} = 15$.

#### Oscillations

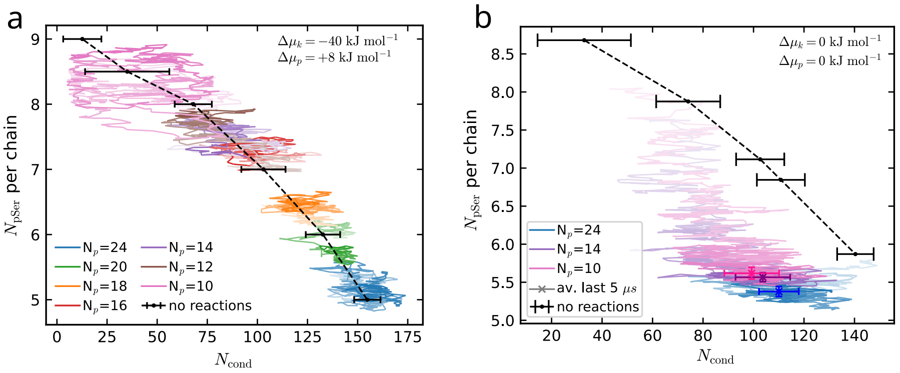


Supplementary Figure 21: Phase portrait with number of chains in condensate on x-axis and number of pSer per chain in y-axis. Black dashed line and points from equilibrium simulations with fixed amount of pSer. (**a**) One example trajectory from active setup with $N_{\rm k} = 2$ and different $N_{\rm p}$, at $\Delta\mu_{\rm k} = -40$ kJ mol$^{-1}$ and $\Delta\mu_{\rm p} = 8$ kJ mol$^{-1}$. (**b**) One example trajectory from passive setup with $N_{\rm k} = 2$ and $N_{\rm p} = 10, 14, 24$, at $\Delta\mu_{\rm k} = 0$ kJ mol$^{-1}$ and $\Delta\mu_{\rm p} = 0$ kJ mol$^{-1}$. Cross markers indicate mean and standard deviation from the last $5{\times}10^8$ MD steps.

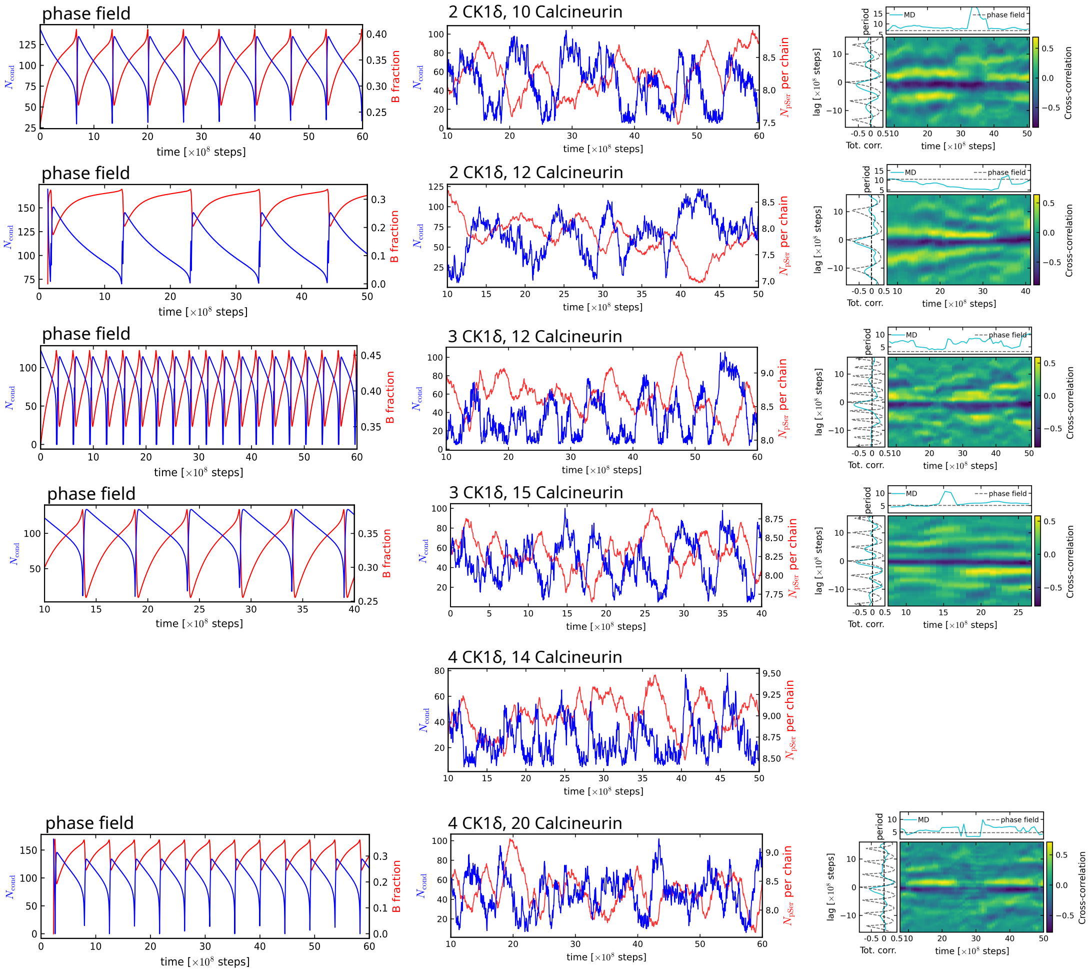


Supplementary Figure 22: On the left, example trajectories of condensate size (blue, left y-axis) and $N_{pSer}$ per chain (red, right y-axis), for each combination of $N_k$ and $N_p$ that show condensation-dissolution oscillating behavior, from MD simulations and phase-field simulation (when available). On the right, the respective cross-correlation between $N_{pSer}$ and condensate size, with sliding windows of 16 $\mu$s. The red curve shows the total cross-correlation from phase-field simulations, when available. On top, the oscillation period from the distance of the 2 central peaks of the cross-correlation with sliding window.

#### Period of oscillations

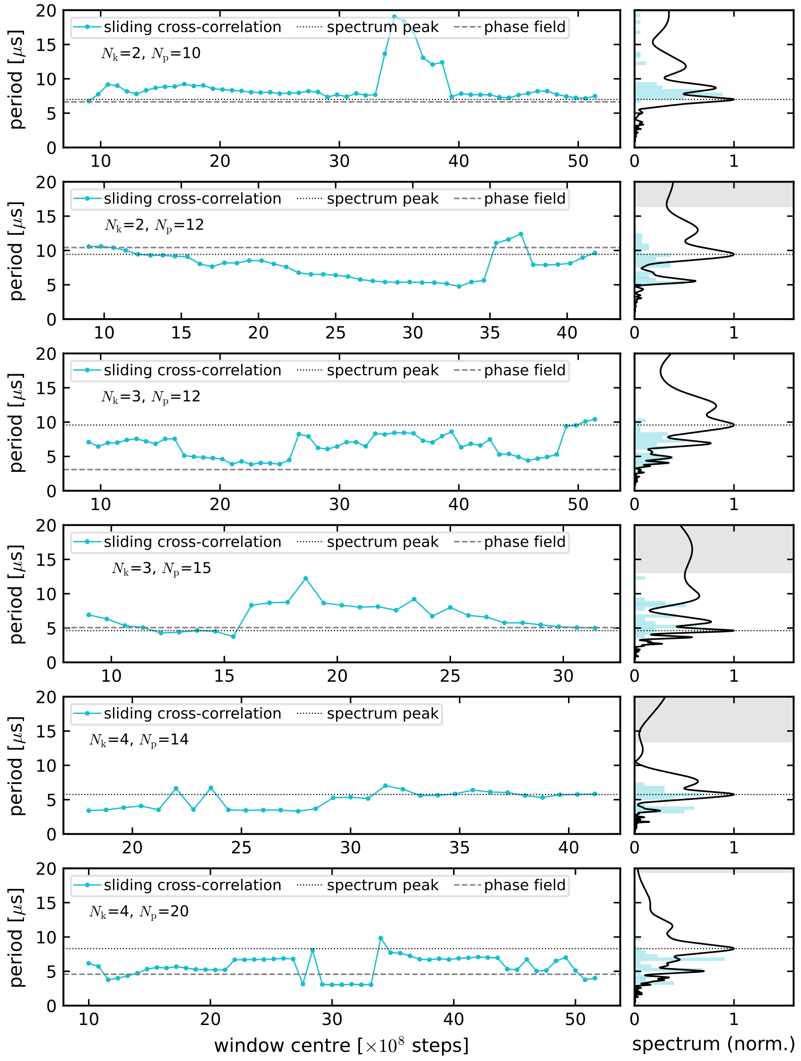


Supplementary Figure 23: On the left, oscillation period from cross-correlation with sliding window of 16 $\mu$s, computed as the interval between the 2 peaks closest to 0 lag (solid cyan line), from peak of the Fourier spectrum of the total cross-correlation (dotted black line) and from phase-field simulations (dashed gray line). On the right, Fourier power spectrum of the detrended total cross-correlation. Shaded bars show the histogram of the sliding-window periods. The gray band marks periods with fewer than three cycles in the trajectory, which the spectrum cannot resolve.

#### Cross-correlations

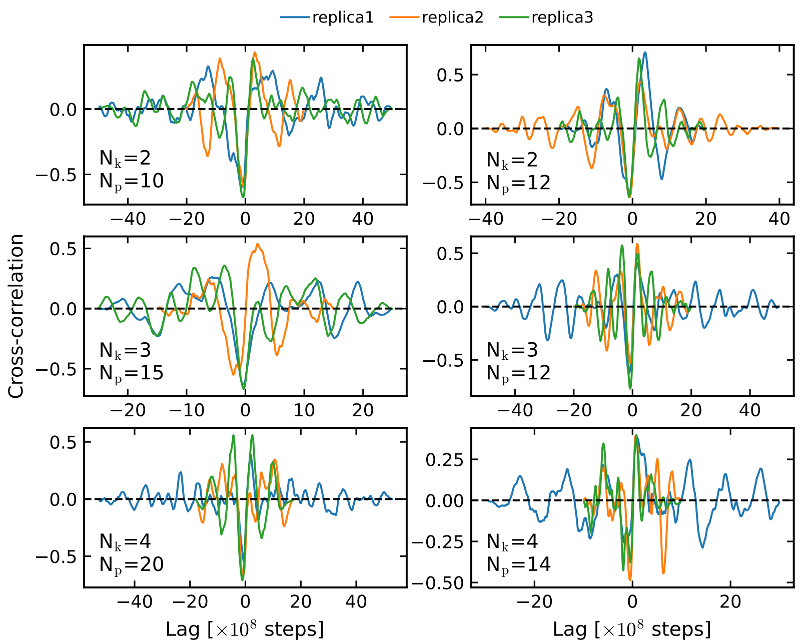


Supplementary Figure 24: Total cross-correlations between number of pSer and condensate size for oscillating systems.

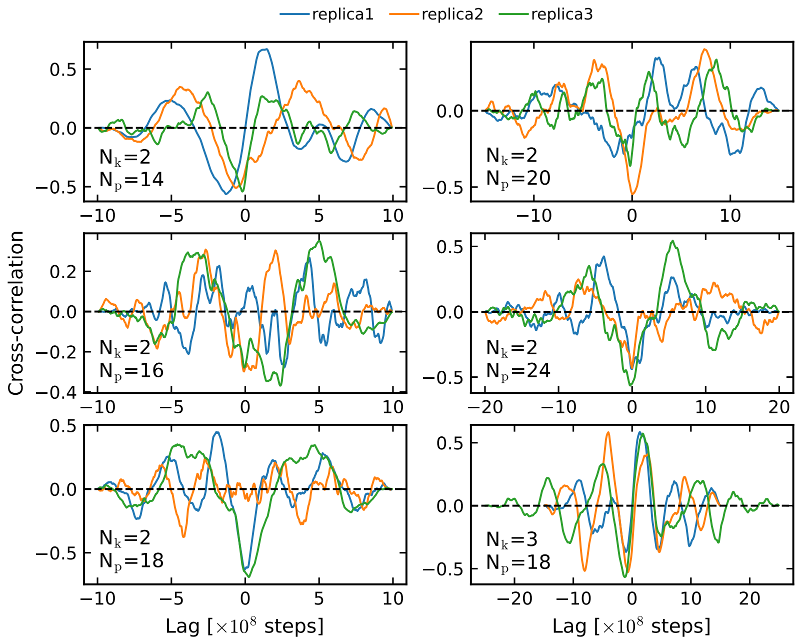


Supplementary Figure 25: Total cross-correlations between number of pSer and condensate size for non-oscillating systems.

#### Relaxation towards fixed point

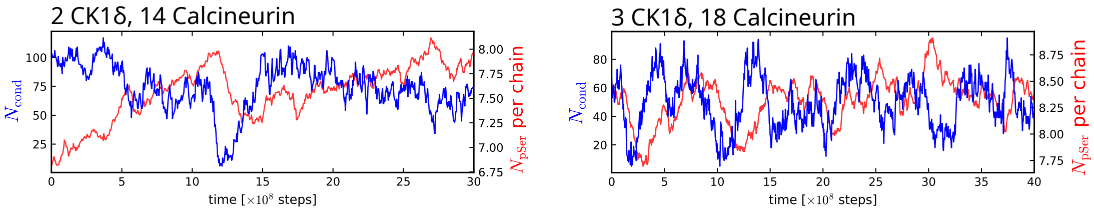


Supplementary Figure 26: Example trajectories of condensate size (blue, left y-axis) and $N_{pSer}$ per chain (red, right y-axis) at $N_k = 2$, $N_p = 14$ and $N_k = 3$, $N_p = 18$.

#### Phase-field model

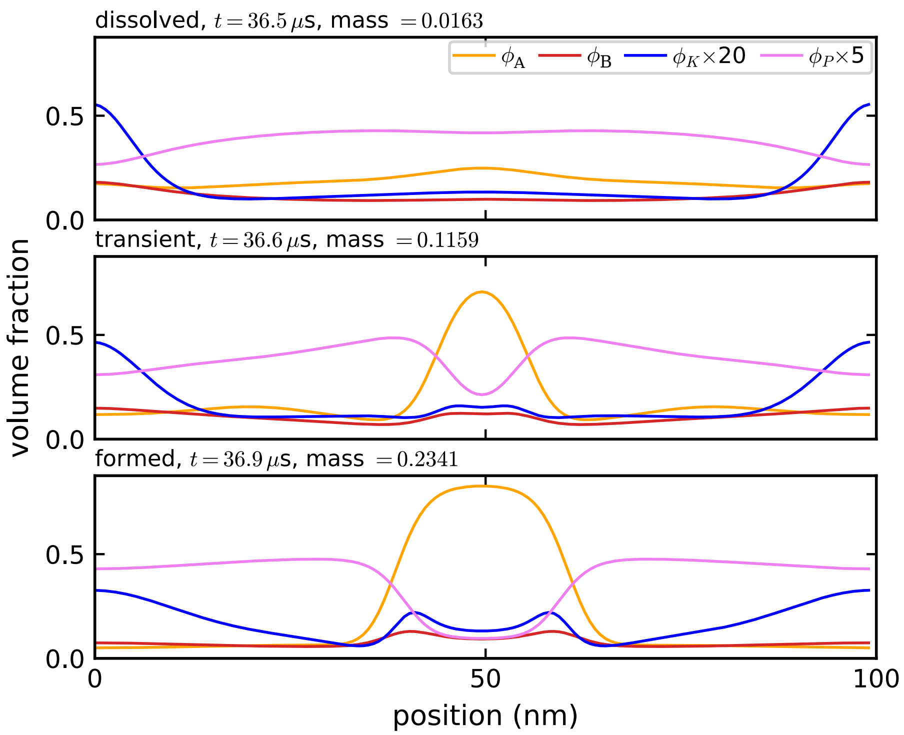


Supplementary Figure 27: Density profiles from phase-field simulation at different times, when condensate is dissolved (top), when it is forming (center), when it is fully formed (bottom). Kinases and phosphatases densities are multiplied by factors 5 and 20 respectively for illustrative purposes.

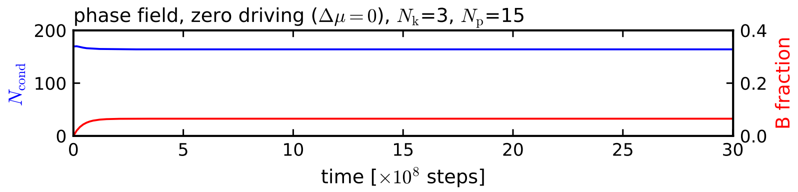


Supplementary Figure 28: Condensate size (blue, left y-axis) and fraction $\phi_B/(\phi_A + \phi_B)$ for phase-field simulations with $N_k = 3$, $N_p = 15$ at zero driving.

#### Independence of limit cycle on initial conditions

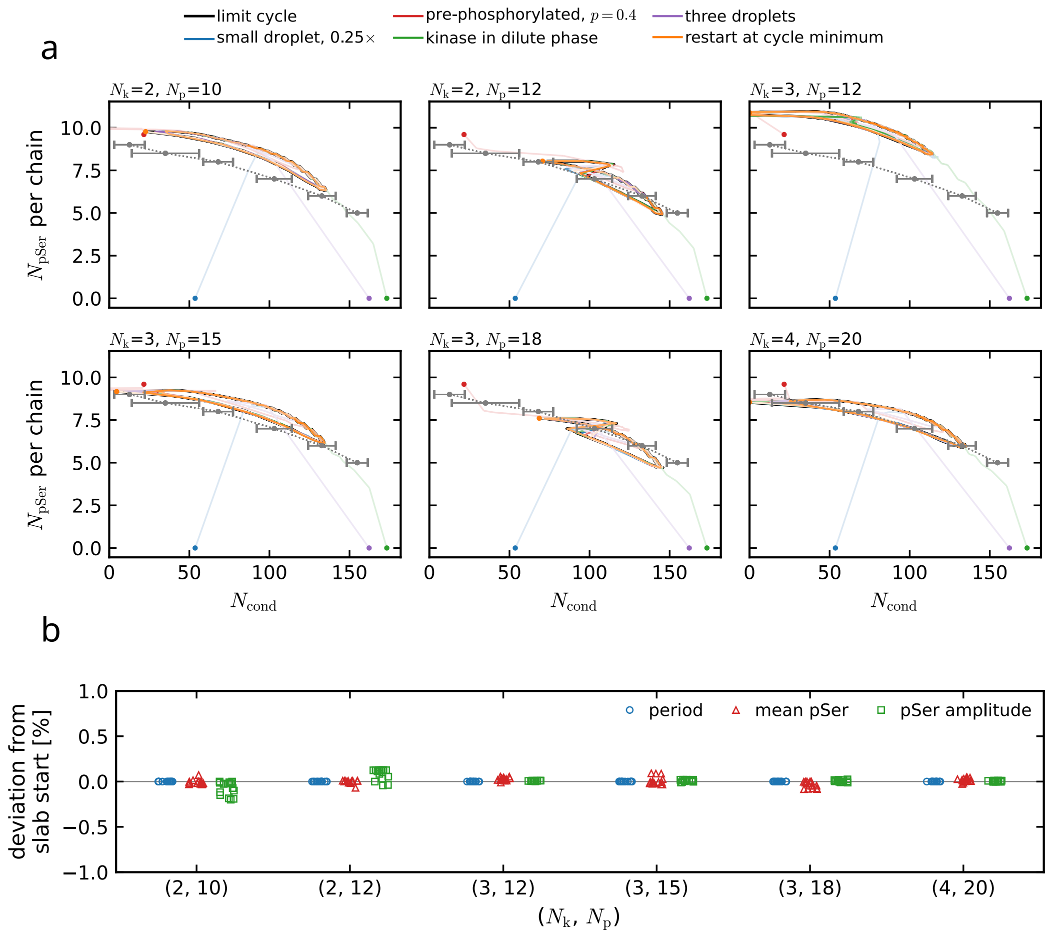


Supplementary Figure 29: (**a**) Phase portraits at the six MD setups $(N_k, N_p)$ where the model oscillates: mean pSer per chain against condensed chains $N_{cond}$ (Eq. **??**), for the first 12 $\mu$s after five different initial conditions (dots): a droplet of a quarter of the coexistence size, a droplet pre-phosphorylated to $p = 0.4$, a droplet with the kinase placed in the dilute phase, three droplets of unequal size, and a restart from the minimum of the cycle. Black dashed line and points from equilibrium simulations with fixed amount of pSer. At every setup all converge onto the limit cycle reached from the condensate slab used for the state diagram (black, last two periods). (**b**) Deviation of the period, the mean pSer per chain and the pSer amplitude from the slab start, for the 15 initial conditions at each of the six setups. Means and amplitudes are taken over a whole number of periods at the end of each 90 $\mu$s run.

#### Relaxation to NESS

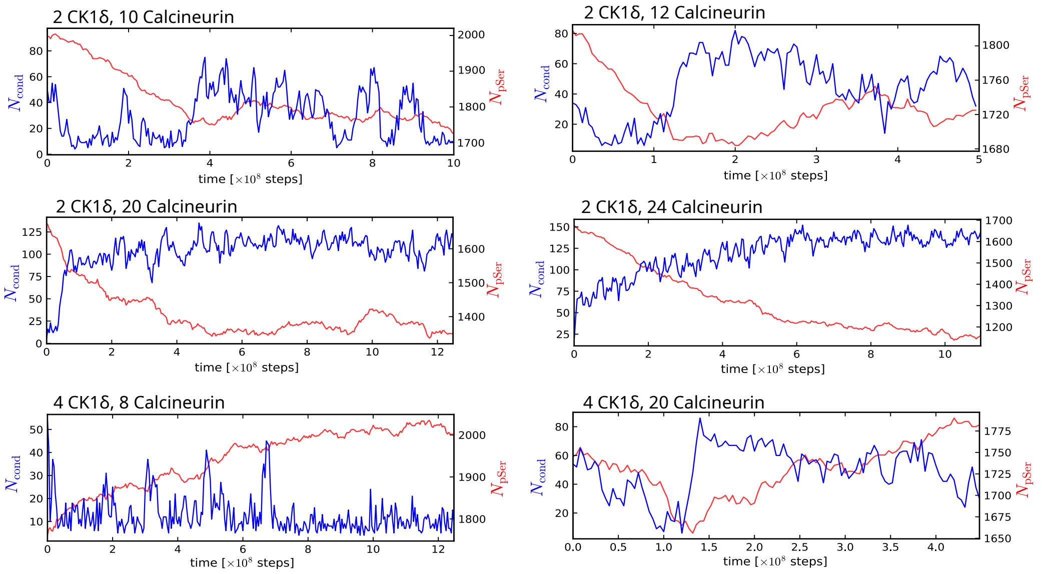


Supplementary Figure 30: Trajectories of condensate size (blue, left y-axis) and $N_{pSer}$ per chain (red, right y-axis) for simulations equilibrating to a NESS, at different amount of kinases and phosphatases.

#### Simulation setup

| **Type** | **TDP-43** | **CK1$\delta$** | **Calcin.** | $(\Delta\mu_k; \Delta\mu_p)$[kJ mol$^{-1}$] | $N_{sim}$:**MD steps** | Therm. |
|---|---|---|---|---|---|---|
| Thermo. check box 30 nm (CALVADOS3) | 1 LCD(Ser395) | - | 1 | (reservoir); -2.5,0,2.5,5 | 5:20×$10^8$ | 50×$10^6$ |
| | 1 5p-LCD(Ser403) | 1 | - | -5,-2.5,0 ; (reservoir) | 5:20×$10^8$ | 50×$10^6$ |
| | 1 5p-LCD(Ser403) | 1 | 1 | -10,-5,0 ; 0 | 5:20×$10^8$ | 50×$10^6$ |
| | 1 5p-LCD(Ser403) | 1 | 1 | -5 ; 2.5 | 5:20×$10^8$ | 50×$10^6$ |
| Dissolution/ condensation/ enzymes position box 100 nm (CALVADOS3) | 200-LCD | 5 | - | -48 ; - | 4:10×$10^8$ | - |
| | 200-LCD(2100pSer) | - | 5 | - ; 8 | 4:20×$10^8$ | - |
| | 200-LCD | 5 | - | - ; - | 1:5×$10^8$ | 100×$10^6$ |
| | 200-LCD | - | 5 | - ; - | 1:5×$10^8$ | 100×$10^6$ |
| | 200-LCD(1257pSer) | 5 | - | - ; - | 1:5×$10^8$ | 100×$10^6$ |
| | 200-LCD(1257pSer) | - | 5 | - ; - | 1:5×$10^8$ | 100×$10^6$ |
| Equilibrium enzymes position box 25×25×150 $nm^3$ (CALVADOS3 & HPS-Urry) | 400-LCD | 5 | - | - ; - | 1:15×$10^8$ | 100×$10^6$ |
| | 400-LCD | - | 5 | - ; - | 1:15×$10^8$ | 100×$10^6$ |
| | 400-LCD(1715pSer) | 5 | - | - ; - | 1:15×$10^8$ | 100×$10^6$ |
| | 400-LCD(1715pSer) | - | 5 | - ; - | 1:15×$10^8$ | 100×$10^6$ |
| | 150-full | 5 | - | - ; - | 1:15×$10^8$ | 100×$10^6$ |
| | 150-full | - | 5 | - ; - | 1:15×$10^8$ | 100×$10^6$ |
| | 150-full(922pSer) | 5 | - | - ; - | 1:15×$10^8$ | 100×$10^6$ |
| | 150-full(922pSer) | - | 5 | - ; - | 1:15×$10^8$ | 100×$10^6$ |
| Reaching NESS box 100 nm (CALVADOS3) | 200-LCD(2000pSer) | 2 | 10 | -40 ; 8 | 3:25×$10^8$ | - |
| | 200-LCD(1818pSer) | 2 | 12 | -40 ; 8 | 3:20×$10^8$ | - |
| | 200-LCD(∼1670pSer) | 2 | 20 | -40 ; 8 | 3:12.5×$10^8$ | - |
| | 200-LCD(∼1670pSer) | 2 | 24 | -40 ; 8 | 3:11×$10^8$ | - |
| | 200-LCD(∼1750pSer) | 4 | 8 | -40 ; 8 | 3:12.5×$10^8$ | - |
| | 200-LCD(∼1750pSer) | 4 | 20 | -40 ; 8 | 3:4.5×$10^8$ | - |
| Enzymes competition box 100 nm (CALVADOS3) | 200-LCD | 2 | 10 | -40 ; 8 | 2:60,1:20×$10^8$ | - |
| | 200-LCD | 2 | 12 | -40 ; 8 | 1:50,2:40×$10^8$ | - |
| | 200-LCD | 2 | 14 | -40 ; 8 | 1:30,2:20×$10^8$ | 10×$10^8$ |
| | 200-LCD | 2 | 16 | -40 ; 8 | 3:20×$10^8$ | 10×$10^8$ |
| | 200-LCD | 2 | 18 | -40 ; 8 | 3:20×$10^8$ | 10×$10^8$ |
| | 200-LCD | 2 | 20 | -40 ; 8 | 3:20×$10^8$ | 5×$10^8$ |
| | 200-LCD | 2 | 24 | -40 ; 8 | 3:30×$10^8$ | 5×$10^8$ |
| | 200-LCD | 3 | 8 | -40 ; 8 | 3:30×$10^8$ | 5×$10^8$ |
| | 200-LCD | 3 | 12 | -40 ; 8 | 1:60,2:20×$10^8$ | 5×$10^8$ |
| | 200-LCD | 3 | 15 | -40 ; 8 | 2:40,1:20×$10^8$ | 5×$10^8$ |
| | 200-LCD | 3 | 18 | -40 ; 8 | 1:40,2:20×$10^8$ | 5×$10^8$ |
| | 200-LCD | 4 | 8 | -40 ; 8 | 3:30×$10^8$ | - |
| | 200-LCD | 4 | 14 | -40 ; 8 | 1:50,2:20×$10^8$ | 10×$10^8$ |
| | 200-LCD | 4 | 20 | -40 ; 8 | 1:60,2:20×$10^8$ | 2×$10^8$ |
| | 200-LCD | 2 | 10 | 0 ; 0 | 1:30×$10^8$ | - |
| | 200-LCD | 2 | 14 | 0 ; 0 | 1:30×$10^8$ | - |
| | 200-LCD | 2 | 24 | 0 ; 0 | 3:30×$10^8$ | 10×$10^8$ |
| | 200-LCD(1000pSer) | - | - | - ; - | 3:5×$10^8$ | 1×$10^8$ |
| | 200-LCD(1200pSer) | - | - | - ; - | 3:5×$10^8$ | 1×$10^8$ |
| | 200-LCD(1400pSer) | - | - | - ; - | 3:5×$10^8$ | 1×$10^8$ |
| | 200-LCD(1600pSer) | - | - | - ; - | 3:5×$10^8$ | 1×$10^8$ |
| | 200-LCD(1700pSer) | - | - | - ; - | 3:5×$10^8$ | 1×$10^8$ |
| | 200-LCD(1800pSer) | - | - | - ; - | 3:5×$10^8$ | 1×$10^8$ |
| 6-state MSM 50k dump | 200-LCD | 2 | 24 | -40 ; 8 | 3:10×$10^8$ | - |
| Phase diagram box 22×22×200 $nm^3$ (CALVADOS3) | 200-LCD(5pSer×ch.) | - | - | - ; - | 3:5×$10^8$ | 1×$10^8$ |
| | 200-LCD(6pSer×ch.) | - | - | - ; - | 3:5×$10^8$ | 1×$10^8$ |
| | 200-LCD(7pSer×ch.) | - | - | - ; - | 3:5×$10^8$ | 1×$10^8$ |
| | 200-LCD(8pSer×ch.) | - | - | - ; - | 3:5×$10^8$ | 1×$10^8$ |
| | 200-LCD(9pSer×ch.) | - | - | - ; - | 3:5×$10^8$ | 1×$10^8$ |
| | 200-LCD(10pSer×ch.) | - | - | - ; - | 3:5×$10^8$ | 1×$10^8$ |
| | 200-LCD(11pSer×ch.) | - | - | - ; - | 3:5×$10^8$ | 1×$10^8$ |

Supplementary Table 1: Summary of simulation setups

#### Supplementary Movies

All simulations show 1 every 5 frames of the original simulation trajectories.

##### Supplementary Movie 1

The movie shows a complete phosphorylation-dephosphorylation cycle from thermodynamic consistency simulation of single 5p TDP-43 LCD chain, single CK1$\delta$ folded domain (blue) and calcineurin (magenta) in cubic box of 30 nm side length. In this simulation, the only phosphosite is Ser 403 (red when phosphorylated). Phosphorylation reactions by CK1$\delta$ are driven with $\Delta\mu_{\mathrm{k}} = -5$ kJ mol$^{-1}$, while dephosphorylation reactions by calcineurin are driven with $\Delta\mu_{\mathrm{p}} = +2.5$ kJ mol$^{-1}$.

##### Supplementary Movie 2

The movie shows the re-condensation of a TDP-43 LCD condensate through enzymatic dephosphorylation from simulation of 200 phosphorylated TDP-43 LCD chains and 5 calcineurin (magenta) in cubic box of 100 nm side length. Serines are shown in yellow, phosphoserines in red. Biggest cluster in cyan is centered in the box.

##### Supplementary Movie 3

The movie shows the ripening of two TDP-43 LCD droplets with active phosphorylation and dephosphorylation reactions, from simulation of 400 TDP-43 LCD chains, 4 CK1$\delta$ (blue) and 48 calcineurin (magenta) in box of dimensions $100 \times 100 \times 200$ nm$^3$. Serines are shown in yellow, phosphoserines in red. The biggest cluster at each snapshot is in cyan and the second biggest cluster in black.

##### Supplementary Movie 4

The movie shows cycles of dissolution and re-condensation of a TDP-43 LCD condensate with active phosphorylation and dephosphorylation reactions from simulation of 200 TDP-43 LCD chains, 2 CK1$\delta$ (blue) and 10 calcineurin (magenta) in cubic box of 100 nm side length. Serines are shown in yellow, phosphoserines in red. Biggest cluster in cyan is centered in the box.

##### Supplementary Movie 5

The movie shows cycles of dissolution and re-condensation of a TDP-43 LCD condensate with active phosphorylation and dephosphorylation reactions from simulation of 200 TDP-43 LCD chains, 3 CK1$\delta$ (blue) and 15 calcineurin (magenta) in cubic box of 100 nm side length. Serines are shown in yellow, phosphoserines in red. Biggest cluster in cyan is centered in the box.

### References


1. Tesei, G. & Lindorff-Larsen, K. Improved predictions of phase behaviour of intrinsically disordered proteins by tuning the interaction range [version 2; peer review: 2 approved]. *Open Research Europe* **2** (2023).
2. Cao, F., von Bülow, S., Tesei, G. & Lindorff-Larsen, K. A coarse-grained model for disordered and multi-domain proteins. *Protein Science* **33,** e5172 (2024).
3. Zippo, E., Dormann, D., Speck, T. & Stelzl, L. S. Molecular simulations of enzymatic phosphorylation of disordered proteins and their condensates. en. *Nat Commun* **16,** 4649 (2025).
4. Mardt, A., Pasquali, L., Wu, H. & Noé, F. VAMPnets for deep learning of molecular kinetics. *Nature communications* **9,** 5 (2018).
5. Dignon, G. L., Zheng, W., Kim, Y. C., Best, R. B. & Mittal, J. Sequence determinants of protein phase behavior from a coarse-grained model. *PLoS Comput. Biol.* **14,** e1005941 (2018).
6. Regy, R. M., Thompson, J., Kim, Y. C. & Mittal, J. Improved coarse-grained model for studying sequence dependent phase separation of disordered proteins. *Protein Science* **30,** 1371–1379 (2021).
7. Gemeinhardt, T. M. *et al.* A disordered linker in the Polycomb protein Polyhomeotic tunes phase separation and oligomerization. *Molecular cell* **85,** 2128–2146 (2025).
